# Lexical semantic content, not syntactic structure, is the main contributor to ANN-brain similarity of fMRI responses in the language network

**DOI:** 10.1101/2023.05.05.539646

**Authors:** Carina Kauf, Greta Tuckute, Roger Levy, Jacob Andreas, Evelina Fedorenko

## Abstract

Representations from artificial neural network (ANN) language models have been shown to predict human brain activity in the language network. To understand what aspects of linguistic stimuli contribute to ANN-to-brain similarity, we used an fMRI dataset of responses to n=627 naturalistic English sentences (Pereira et al., 2018) and systematically manipulated the stimuli for which ANN representations were extracted. In particular, we i) perturbed sentences’ word order, ii) removed different subsets of words, or iii) replaced sentences with other sentences of varying semantic similarity. We found that the lexical semantic content of the sentence (largely carried by content words) rather than the sentence’s syntactic form (conveyed via word order or function words) is primarily responsible for the ANN-to-brain similarity. In follow-up analyses, we found that perturbation manipulations that adversely affect brain predictivity also lead to more divergent representations in the ANN’s embedding space and decrease the ANN’s ability to predict upcoming tokens in those stimuli. Further, results are robust to whether the mapping model is trained on intact or perturbed stimuli, and whether the ANN sentence representations are conditioned on the same linguistic context that humans saw. The critical result—that lexical-semantic content is the main contributor to the similarity between ANN representations and neural ones—aligns with the idea that the goal of the human language system is to extract meaning from linguistic strings. Finally, this work highlights the strength of systematic experimental manipulations for evaluating how close we are to accurate and generalizable models of the human language network.

## 1. Introduction

Research in psycholinguistics and cognitive neuroscience of language strives to understand the representations and algorithms that support human comprehension and production abilities. Until recently, mechanistic accounts of human language processing have been out of reach. However, artificial neural network (ANN) language models now hold substantial promise for developing and evaluating computationally precise hypotheses about language processing. In particular, contemporary ANN language models achieve impressive performance on a variety of linguistic tasks (e.g., Devlin et al., 2018; Liu et al., 2019; Brown et al., 2020; Rae et al., 2021; Chowdhery et al., 2022; OpenAI, 2023). Furthermore, representations extracted from ANN language models—especially unidirectional attention Transformer architectures like GPT2 (Radford et al., 2019) can explain substantial variance in brain activity recorded from the human language network using regression-based evaluation metrics (e.g., Jain & Huth, 2018; Gauthier & Levy, 2019; Toneva & Wehbe, 2019; Schrimpf et al., 2021; Caucheteux & King, 2022; Goldstein et al., 2022; Hosseini et al., 2022; Kumar et al., 2022; Oota et al., 2022; Pasquiou et al., 2022). This correspondence has been suggested to derive, at least in part, from the convergence of the ANNs’ linguistic representations with those in the human brain (Schrimpf et al., 2021; Caucheteux & King, 2022; Goldstein et al., 2022; Hosseini et al., 2022), despite the vast differences in their learning and architecture (e.g., Huebner & Willits, 2021; Warstadt & Bowman, 2022).

However, many questions remain about the factors that contribute to ANN-to-brain mapping, i.e., the ability to predict brain responses from ANN representations. One critical question concerns the *aspects of linguistic content and form* that play a role. To shed light on this question, we used a published fMRI dataset (Pereira et al., 2018) where brain responses were collected from ten native speakers of English as they read syntactically and semantically diverse passages, consisting of several sentences each. We reproduced the result of the top-performing brain-encoding ANN language model in Schrimpf et al. (2021)—GPT2-xl (Radford et al., 2019)—on this dataset, and investigated what drives the model’s brain predictivity (or ‘brain score’; Schrimpf et al., 2018). In particular, we evaluated the contributions to accurate mapping of *sentence meaning* (largely carried by content words) and *syntactic form* (conveyed via word order and function words), along with superficial control features, like sentence length. To do so, we performed 12 sets of experiments: 3 categories of linguistic manipulations across 4 variants of what we term here ‘computational experimental design’, as elaborated next.

First, we systematically manipulated the linguistic stimuli in 3 ways: by altering the word order of the sentence (across 7 conditions; see <u>Methods; Perturbation manipulation conditions</u> for details), omitting different subsets of words (across 5 conditions), and replacing a sentence with sentences of different degrees of semantic relatedness (across 4 conditions). Some of these manipulations disrupt the syntactic form of the sentence (e.g., changing the order of the words or removing the function words); whereas other manipulations affect sentence meaning (e.g., removing the content words or replacing a sentence with a semantically unrelated sentence). We asked how well ANN representations for the resulting altered stimuli (across the 16 conditions) can predict neural responses compared to the ANN representations of a) the original, unaltered sentence and b) a control, length-matched condition (a random list of words). Next, we explored possible causes for the differential effects of these manipulations on brain predictivity by examining the changes in the ANN representations and next-word prediction task performance as a function of stimulus alterations. Finally, to evaluate the robustness of the results to the computational experiment design, we performed all three types of linguistic manipulations across four experimental set-ups, crossing i) whether the model that maps from stimuli to brain representations was trained on intact or perturbed stimuli; and ii) whether the ANN representations of the target sentences were contextualized with respect to the preceding sentences in a passage or not.

### Manipulations of linguistic stimuli

Language allows its users to package meanings into sequences of words. Content words, like nouns, verbs and adjectives (which carry the most information in a sentence, as can be quantified using information-theoretic measures; e.g., Shannon, 1948) have to be selected, so as to capture the right word-level meanings, and then they have to be assembled into phrases and sentences according to the rules of the language. The assembly process includes ordering the words in a particular way as well as adding appropriate inflectional morphological markers and function words, like determiners and prepositions. The result of all these operations is a meaningful and well-formed sentence. Here we evaluate the relative importance of different aspects of linguistic stimuli (sentences) for ANN-to-brain mapping. To do so, we systematically manipulate the sentences across three manipulation categories (**Figure 1A; Table 1**) before passing them into the ANN and use the resulting ANN model representations to predict brain responses (**Figure 1B**).

**Figure 1.**
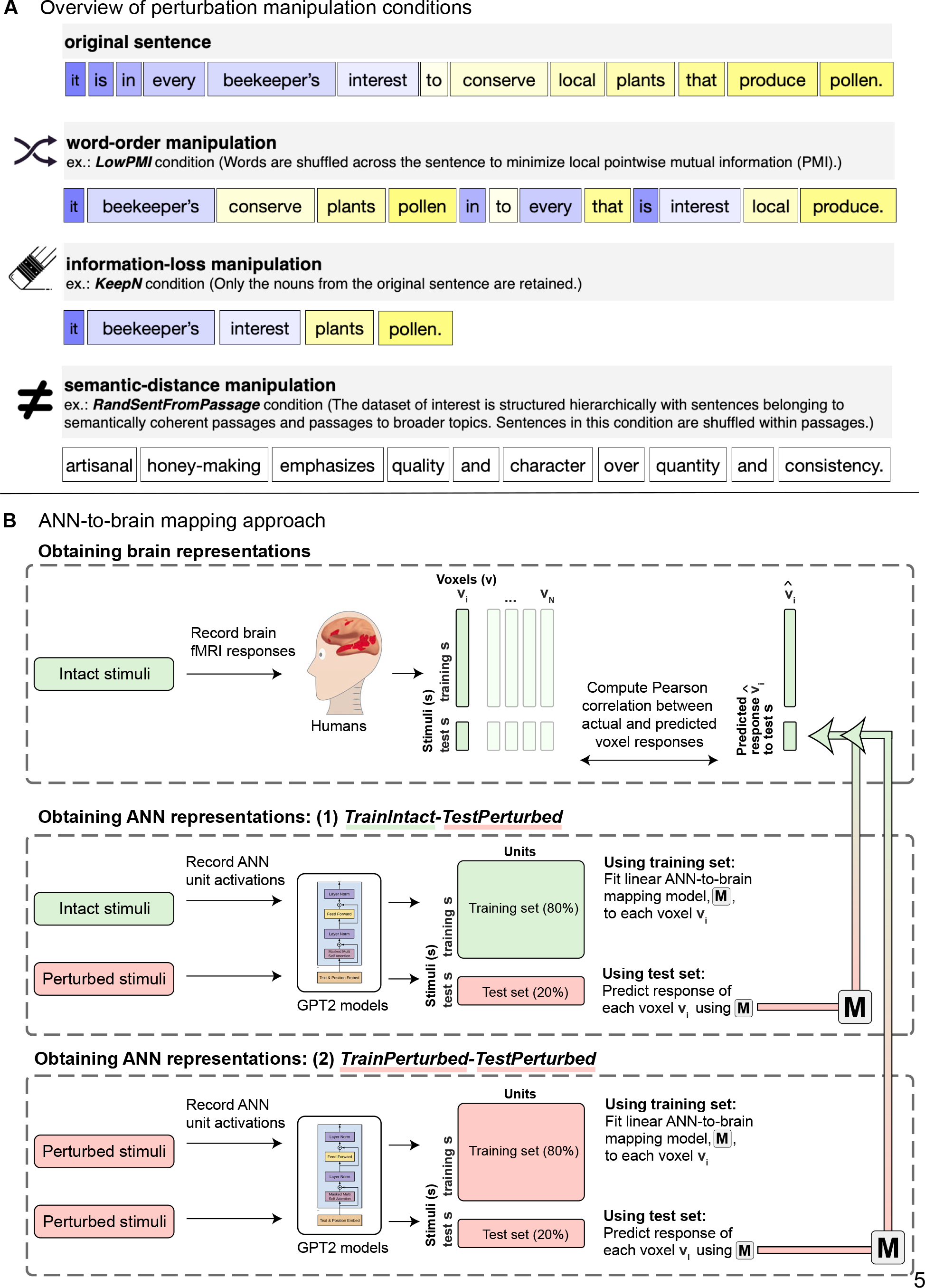
Overview of perturbation manipulation conditions and the ANN-to-brain mapping approach. **A.** Overview of perturbation manipulation conditions. An example (original) sentence is illustrated together with an example of the stimulus in a sample condition from each of the three types of perturbation manipulations (word-order manipulations, information-loss manipulations, and semantic-distance manipulations), as detailed in <u>Methods;</u> <u>Perturbation manipulation conditions</u>. **B.** Overview of the ANN-to-brain mapping approach. Brain data from human participants (n=10) were recorded while they read intact sentences using functional magnetic resonance imaging (fMRI) (Pereira et al., 2018). Brain data consisted of voxel responses within the language-selective network (individually defined using an independent localizer task; Fedorenko et al., 2010) for each of the 10 participants. Following Schrimpf et al. (2021), we divided the stimuli (i.e., sentences) into training/test sets. We then retrieved ANN model representations for the stimuli and fitted a linear ANN-to-brain mapping model (**M**) from the ANN representations of the training stimuli to each single voxel’s (within the language network) corresponding recordings for those stimuli (the fitting process is not illustrated in this graphic). Next, we tested the ANN-to-brain mapping model (**M**) on the ANN representations of the held-out test stimuli to generate predicted brain responses for those stimuli, for each voxel (illustrated by the gradient arrows). Lastly, we compared predicted versus actual brain responses for each voxel using the Pearson correlation coefficient. This process was repeated five times, holding out a different set of 20% of stimuli each time. ANN representations were obtained using two approaches (for the motivation and details, see <u>Methods;</u> <u>Manipulations of computational experimental design</u>): **(1) *TrainIntact-TestPerturbed***, where ANN representations for the training set were obtained from the original, intact stimuli, whereas the ANN representations for the test set were obtained from the perturbed stimuli; and **(2) *TrainPerturbed-TestPerturbed***, where ANN representations for the training *and* test set were obtained from the perturbed stimuli. These two approaches were crossed with whether the preceding sentences in the passage were included as contextualizing input for the ANN (not depicted). *The GPT2 illustration was adapted from Radford et al. (2018)*.

**Table 1.** Overview of perturbation manipulation conditions and sample stimuli for each condition. The perturbation manipulation conditions that we use in the current work are motivated by prior theorizing in language research and/or past empirical findings from both neuroscience and natural language processing (NLP). The perturbation manipulations include i) word-order manipulations of varying severity that preserve or destroy local dependency structure (following Mollica, Siegelman et al., 2020) allowing us to investigate the effect of word order degradation while controlling for local word co-occurrence statistics; ii) information-loss manipulations with deletion of words of different parts of speech (following O’Connor & Andreas, 2021) allowing us to investigate loss of information from particular classes of words; iii) semantic-distance manipulations with sentence substitutions that relate to the meaning of the original sentence to varying degrees (inspired by Pereira et al., 2018) allowing us to investigate loss of semantic and more general topical information while retaining sentence well-formedness. Lastly, as a baseline length-matched control condition, we include a random word list, where each word is substituted with a different random word.

| Manipulation type | Condition name | Sample stimulus |
| --- | --- | --- |
| **original** | *Original* | it is in every beekeeper's interest to conserve local plants that produce pollen. |
| **word-order** | *1LocalWordSwap* | it is in every beekeeper's interest to conserve local plants produce that pollen. |
|  | *3LocalWordSwaps* | in it is every beekeeper's interest to conserve local plants produce that pollen. |
|  | *5LocalWordSwaps* | in it is every interest beekeeper's to conserve local plants produce pollen that. |
|  | *7LocalWordSwaps* | in every it is interest beekeeper's to conserve plants local that pollen produce. |
|  | *ReverseOrder* | pollen produce that plants local conserve to interest beekeeper's every in is it. |
|  | *LowPMI* | it beekeeper's conserve plants pollen in to every that is interest local produce. |
|  | *LowPMIRand* | in that pollen to is plants every beekeeper's conserve produce interest it local. |
| **information-loss** | *KeepContentW* | it is beekeeper's interest conserve local plants produce pollen. |
|  | *KeepNVAAdj* | it is beekeeper's interest conserve local plants produce pollen. |
|  | *KeepNV* | it is beekeeper's interest conserve plants produce pollen. |
|  | *KeepN* | it beekeeper's interest plants pollen. |
|  | *KeepFunctionW* | in every to that. |
| **semantic-distance** | *Paraphrase* | conserving regional vegetation that provides pollen is of the utmost importance for beekeepers. |
|  | *RandSentFromPassage* | beekeepers also discourage the use of pesticides on crops because they could kill the honeybees. |
|  | *RandSentFromTopic* | artisanal honey-making emphasizes quality and character over quantity and consistency. |
|  | *RandSent* | mosquitos are thin small flying insects that emit a high-pitched sound. |
| **control** | *RandWordList* | of shears metallic is in individual machine for fracture a singer can have. |

#### i) Word-order manipulations

The first class of manipulations targets the **order of the words** in the sentence. Word order is an important cue to how words relate during sentence comprehension (e.g., Bever, 1970; Kimball, 1973). However, word order rigidity varies across languages, with some languages exhibiting flexible orderings, pointing to a more limited role of word order, at least in those languages (e.g., Hale, 1983; Dryer & Haspelmath, 2013; Jackendoff & Wittenberg, 2014). Moreover, work in psycholinguistics has shown that comprehension is highly robust to errors in the linguistic input, including word order errors, as long as a plausible meaning can be recovered. For example, given the sentence *The mother gave the candle the daughter*, people typically infer the intended meaning to be the more plausible *The mother gave the daughter the candle*, suggesting that word order information can be overridden in favor of a plausible meaning (e.g., Levy et al., 2009; Gibson et al., 2013). Furthermore, word-order transpositions, as in *You that read wrong again!*, often go unnoticed during sentence reading (Mirault et al., 2018; Wen et al., 2021).

Similarly, recent evidence from neuroscience has shown that the human language network—a set of brain areas that are selectively and robustly activated when humans process language (e.g., Fedorenko et al., 2011; Regev et al., 2013; Fedorenko & Thompson-Schill, 2014; Lipkin et al., 2022)—exhibits the same amount of activation to word-order-manipulated sentences as to intact ones as long as the pointwise mutual information among nearby words remains as high as in the intact sentences (which presumably allows for the formation of local semantic and syntactic dependencies) (Mollica, Siegelman et al., 2020). This result aligns with findings by Gauthier & Levy (2019), who showed that fine-tuning the pre-trained BERT language model (Devlin et al., 2018) on a word prediction task that selects against word-order-based representations of the input (namely, fine-tuning on a corpus where words were randomly shuffled within a sentence) leads to an increase in brain decoding performance for the same fMRI benchmark used here (Pereira et al., 2018). Lastly, recent work in natural language processing (NLP) has shown that word-order information is not necessarily needed to solve many current NLP benchmark tasks (e.g., Pham et al., 2021; Sinha et al., 2021; Papadimitriou et al., 2022; cf. Abdou et al., 2022; Lasri et al., 2022), although these results may be more reflective of how the benchmarks were constructed rather than something about human language or language-processing mechanisms (e.g., McCoy et al., 2019).

Therefore, given this evidence from NLP and language neuroscience, we hypothesized that ANN-to-brain mapping performance on current fMRI datasets may similarly not require ANNs to leverage word-order information. Building on Mollica, Siegelman et al. (2020), we evaluate both local scrambling of words (which may better preserve syntactic and semantic dependencies among nearby words) and more global re-arrangement of words (which does not preserve such dependencies).

#### ii) Information-loss manipulations

The second class of manipulations targets the information carried by **words from specific parts of speech**. Research in formal semantics has traditionally posited a categorical difference between the semantics of content (open-class/lexical) words and function (closed-class/logical) words, whereby function words, unlike content words, are assumed to carry little lexical meaning and primarily encode logical relationships between content words (e.g., Partee, 1992; Chierchia, 2013). Further, research in distributional semantics suggests that the semantic properties of function words are not well-represented by word co-occurrence distributions, in sharp contrast with the meanings of content words, which are better captured by distributional patterns (Boleda, 2020; though see e.g., Baroni et al., 2012; Linzen et al., 2016; Abrusán et al., 2018).

Indeed, previous co-occurrence-based models often benefited from the removal of function words and other high-frequency “stop words” for solving various NLP tasks (e.g., Bernardi et al., 2013; Herbelot & Baroni, 2017; Lazaridou et al., 2017). Even for state-of-the-art ANN language models, it has recently been shown that retaining only the content words in linguistic context has little effect on next-word prediction performance, with performance varying as a function of how much of the lexical content is included (i.e., higher performance when keeping *all* content words vs. keeping only subsets, such as keeping only the nouns and verbs, or keeping only the nouns) (O’Connor & Andreas, 2021). Whereas function words have been shown to have a sizable effect on ANN next-word prediction performance within local sentence contexts (because they help ensure grammaticality) (Khandelwal et al., 2018), content words strongly influence prediction performance both within local and more extended contexts (Khandelwal et al., 2018; O’Connor & Andreas, 2021).

Research from psycholinguistics similarly shows an asymmetry between content and function words: when reading sentences, people tend to overlook the omission or repetition of function words, but such errors are much more noticeable for content words (Staub et al., 2019; Huang & Staub, 2021). Differences can also be found in language production: function words tend to have shorter pronunciations than content words of the same length, because they are typically highly predictable from context, which leads to phonological reduction and de-stressing (e.g., Bell et al., 2009).

In line with this previous research, we hypothesized that removing content words, but not function words, should have a strong negative effect on ANN-to-brain mapping, and that brain predictivity should increase the more lexical content of the original sentence is available to the ANN for building a representation. Following O’Connor & Andreas (2021), we evaluate the effect of preserving all or some of the content words (like nouns and verbs) or function words.

#### iii) Semantic-distance manipulations

The third class of manipulations targets **sentence-level meanings** and serves to test how precisely the meaning of a sentence has to be encoded for a successful ANN-to-brain mapping.

Previous research in computational neuroscience has shown that vectors that represent lexical semantics based on word co-occurrence statistics (GloVe; Pennington et al., 2014) can be decoded from fMRI data recorded while participants read sentences (Pereira et al., 2018). In particular, Pereira et al. (2018) demonstrated that sentence pairwise classification accuracy depended on how semantically similar the sentence pairs were: sentences from different topics (e.g., a sentence about beekeeping vs. a sentence about skiing) were easier to distinguish than sentences that talk about different ideas within the same general topic (e.g., two sentences about beekeeping that come from distinct passages: e.g., one passage about the importance of beekeeping for the health of the planet and the other telling a story about a particular beekeeper), with sentences that talk about related ideas within the same topic (e.g., two sentences from the same passage on the importance of beekeeping) being the most challenging to distinguish.

However, in Pereira et al. (2018), even neural responses to sentences from the same passage could still be reliably discriminated, suggesting that semantic representations of linguistic input are relatively fine-grained in both fMRI data and word embeddings. On the other hand, even neural responses to sentences from distinct topics could not be *perfectly* discriminated (classification accuracy: 81-84%, chance level: 50%), suggesting that representations of sentences that are unrelated to the target sentence may share some features with the representation of the target sentence that can lead to misclassification (at least when using coarse representations based on averaged decontextualized GloVe embeddings for decoding). This pattern of results raises the question of how much of the fMRI signal that ANN-to-brain models are able to predict represents a sentence’s exact or approximate semantic content.

We manipulate sentence meaning by replacing the original sentence with sentences that vary in how similar they are in meaning to the original. In line with Pereira et al. (2018), we hypothesized that substituting sentences that are semantically more distant from the target sentence would elicit lower ANN-to-brain mapping performance. Further, because ANN representations of compositional sentence meaning (as investigated here) are richer and finer-grained than a simple average of decontextualized word embeddings, we hypothesized that the representations of topically unrelated sentences would be more distant, so that substituting them would result in very low brain predictivity. We leveraged the hierarchical structure of the linguistic materials in the Pereira et al. (2018) fMRI benchmark to vary semantic distance between the original sentences and the manipulated ones, in a similar way to the original study, and additionally created paraphrases for each sentence, which were expected to elicit high ANN-to-brain performance.

### Understanding the effects of linguistic manipulations on brain predictivity

To complement our findings across the linguistic perturbation manipulations, we present a set of exploratory analyses that aim to uncover potential causes for the differential effects of perturbation manipulations on brain predictivity. In particular, we look for possible correlates of brain predictivity across perturbation manipulation conditions in a) the **ANN’s representational space** and b) the **ANN’s performance on the next-word prediction task** for the manipulated sentence sets.

The investigation of the **ANN representational space** is motivated by the use of representational similarity metrics to compare high-dimensional vectors derived from brain recordings, ANNs, or both (Kriegeskorte et al., 2008; Kriegeskorte, 2015). Representational Similarity Analysis (RSA) relies on the strength of correlation between sets of vectors to make inferences about the information contained in the distributed patterns within the vectors. Although originally developed to compare vectors derived from different brain recording modalities, RSA-style approaches can also be used to compare representations derived from various instantiations of ANNs (e.g., Morcos et al., 2018; Barrett et al., 2019; Kornblith et al., 2019). Here, we investigated how the ANN representational space is transformed by different perturbation manipulations. We hypothesized that larger transformations in the ANN representational space (relative to the representational space for the original, intact stimuli) would be associated with larger changes in brain predictivity.

The investigation of **ANN task performance** is motivated by research in psycholinguistics and neuroscience suggesting that predictive processing is a core mechanism for language comprehension (*within psycholinguistics:* Rayner et al., 2006; Demberg & Keller, 2008; Bicknell et al., 2010; Smith & Levy, 2013; Brothers & Kuperberg, 2021; *within neuroscience*: Henderson et al., 2016; Willems et al., 2016; Lopopolo et al., 2017; Heilbron et al., 2019; Shain, Blank et al., 2020). Converging research from computational neuroscience has reported a positive correlation between the next-word prediction performance of ANN language models and model-to-brain correspondence (Schrimpf et al., 2021; Caucheteux & King, 2022; Goldstein et al., 2022; Hosseini et al., 2022; cf. Antonello & Huth, 2022). In line with this evidence, we hypothesized that ANN brain predictivity would correlate with how well the model can predict upcoming words within a manipulated sentence, such that manipulations that render sentences less predictable would, on average, lead to representations that map less well onto human brain data.

### Manipulations of computational experimental design

ANN-based modeling of language provides a novel toolkit for testing theoretically-motivated hypotheses about language processing in the mind and brain. However, results derived via such in-silico investigations of human language processing may not be robust to variations in *how* the ANN-to-brain match comparisons are performed. Here, we evaluate the relative importance of manipulations to what we denote as the ‘computational experimental design’ for ANN-to-brain match performance to evaluate the robustness of our results. Specifically, we investigate the contribution of two factors, as elaborated below: i) whether the training data stimuli for the ANN-to-brain mapping model are intact or perturbed, and ii) whether the target sentence is contextualized with preceding sentences from the passage.

#### i) ANN-to-brain mapping model training stimuli

In our main approach (*TrainIntact-TestPerturbed,* as illustrated in **Figure 1B**, upper two panels), we train a linear ANN-to-brain mapping model using ANN representations for the original (intact) stimuli from Pereira et al. (2018) and human brain responses obtained during the processing of the same, intact versions of the stimuli. This training set-up corresponds to the main use case of ANN-to-brain encoding models and follows prior work (e.g., Schrimpf et al., 2021; Caucheteux & King, 2022; Goldstein et al., 2022). Using this standard set-up, we investigated our main research question, i.e., *which aspects of a linguistic stimulus contribute to successful ANN-to-brain mapping performance*, by evaluating the mapping model’s ability to predict brain responses to intact sentences from ANN representations of sentences that were perturbed in one of the ways described above (detailed description in <u>Methods; Perturbation manipulation conditions</u>). If degrading a particular aspect of the sentence decreases the mapping model’s performance (brain predictivity), we would like to conclude that this aspect of the stimulus is a critical contributor to the mapping model’s performance. If, on the other hand, degrading the stimulus does not lead to lower brain predictivity, we would like to conclude that the mapping model does not pay any appreciable attention to the part of the ANN representation of the stimulus that is sensitive to the removed information.

However, there are (at least) two possible explanations for why sentence perturbation manipulations may adversely affect the success of an ANN-to-brain mapping model trained on representations of intact sentences: either a successful mapping must critically rely on the information removed by a given perturbation (our desired interpretation, as stated above), or perturbing the input to the ANN model only at test time introduces out-of-distribution inputs to the trained ANN-to-brain mapping model, i.e., lower brain predictivity can be explained by a distribution shift of the input to the mapping model at test time. To distinguish between these possibilities, we tested how well an ANN-to-brain mapping model can predict brain responses when the input to the ANN is perturbed in the same way during training and testing (**Figure 1B**, *TrainPerturbed-TestPerturbed*). When the ANN-to-brain mapping models are trained *and* tested on the same set of perturbations, a decline in brain predictivity relative to the performance using intact sentences (*Original* benchmark) cannot be explained by a distribution shift in the input to the model at test time. Thus, this approach can reveal the degree to which perturbations remove information from the ANN representation of the stimuli that is useful for a mapping model when it learns a relationship between ANN representations and brain data.

#### ii) Contextualization of the linguistic stimuli

ANN-to-brain predictivity analyses typically aim to mimic the experimental procedure for which the human brain data was obtained. Linguistic stimuli in human brain imaging studies are often contextualized within a story (e.g., Blank et al., 2014; Huth et al., 2016; Schoffelen et al., 2019) or a passage (e.g., Pereira et al., 2018). Given that large-scale ANN language models (such as Transformer language models) are able to condition input representations on large amounts of preceding linguistic context, they enable mimicking the human experimental design by providing the same linguistic stimuli as context to the ANN as were provided to the human participants. However, whether this is the right approach, empirically and conceptually, is not clear.

On the one hand, providing the same context to the ANN language models for representation building as what humans saw/heard during the experiment could improve ANN-to-brain mapping performance by modulating sentence representations in ways similar to how the human brain is affected by context. On the other hand, sentence contextualization could hurt match-to-brain performance. First, the way in which ANNs vs. humans represent contextual information in memory is likely very different. In particular, constrained by memory limitations, humans do not retain detailed linguistic representations of the preceding context (e.g., Potter et al., 1980; Potter & Lombardi, 1990; Potter, 2012; Futrell et al., 2020); instead, as they process linguistic input, they appear to extract the representations of the relevant *meaning* and ‘discard’ the exact word sequences (e.g., Christiansen & Chater, 2016; Potter & Lombardi, 1998). And second, human neuroscience studies have suggested that extended story contexts are represented not in the language network proper (which we focus on here), but in a distinct brain network—the Default network (e.g., Lerner et al., 2011; Simony et al., 2016; I. A. Blank & Fedorenko, 2020). Thus, neural responses to language stimuli of the language-selective areas may only be capturing the local processing of the current sentence and would therefore align better with decontextualized sentence representations (see Jain & Huth, 2018; Caucheteux et al., 2021 for evidence that ANN representations with varying amounts of linguistic context lead to differential mapping performance with different brain areas). We therefore evaluated brain predictivity for sentence representations with and without contextualization through inclusion of preceding sentences in the passage; we refer to these as *contextualized* and *decontextualized* sentence representations respectively. We did this for the two ANN-to-brain mapping model training approaches introduced above (i.e., *TrainIntact-TestPerturbed* and *TrainPerturbed-TestPerturbed*).

To foreshadow our results, we find that i) lexical-semantic content of the sentence, rather than syntactic structure (conveyed via word order or function words), is responsible for the ability of ANNs to predict fMRI responses in the human language network. We further show that ii) linguistic perturbations that decrease brain predictivity have interpretable causes: they lead to a) more divergent representations in the ANN’s embedding space (relative to the representations of intact sentences) and b) a decrease in the ANN’s next-word prediction task performance, i.e., its ability to predict upcoming tokens in those stimuli. Finally, iii) the results from the linguistic manipulations are largely robust to variations in the computational experimental design, which impact the overall magnitude of brain scores but not their pattern across conditions.

## 2. Methods

Here we describe in detail (i) our manipulations of the linguistic stimuli that are designed to isolate the influences of different features of the input, (ii) how we obtain ANN representations for these stimuli, and (iii) how we perform ANN-to-brain mappings. Because our approach is based on the ANN-to-brain mapping framework from Schrimpf et al. (2021), the sections <u>Methods; Comparison</u> <u>of ANN model representations to brain measurements</u>, <u>fMRI dataset (*Pereira2018)*</u>, and <u>Estimation of ceiling</u> are similar to the methods reported in Schrimpf et al. (2021). All code is publicly available at https://github.com/carina-kauf/perturbed-neural-nlp.

### fMRI dataset

We used the data from Pereira et al.’s (2018) Experiments 2 (n=9) and 3 (n=6) (10 unique participants, all native speakers of English). (The set of participants is not identical to Pereira et al., 2018: i) one participant (tested at Princeton) was excluded from both experiments here to keep the fMRI scanner the same across participants; and ii) two participants who were excluded from Experiment 2 in Pereira et al. (2018) based on the decoding results in Experiment 1 of that study were included here, to err on the conservative side.) Stimuli for Experiment 2 consisted of 384 sentences (96 text passages, four sentences each), and stimuli for Experiment 3 consisted of 243 sentences (72 text passages, three or four sentences each). The two sets of materials were constructed independently, and each spanned a broad range of content areas. Sentences were 7-18 words long in Experiment 2, and 5-20 words long in Experiment 3. The sentences were presented on the screen one at a time for 4s each (followed by 4s of fixation, with additional 4s of fixation at the end of each passage), and each participant read each sentence three times, across independent scanning sessions (see Pereira et al., 2018 for details of experimental procedure and data acquisition).

#### Preprocessing and response estimation

Data preprocessing was carried out with SPM5 (using default parameters, unless specified otherwise) and supporting, custom MATLAB scripts. Preprocessing included motion correction (realignment to the mean image of the first functional run using 2nd-degree b-spline interpolation), normalization (estimated for the mean image using trilinear interpolation), resampling into 2 mm isotropic voxels, smoothing with a 4mm FWHM Gaussian filter and high-pass filtering at 200s. A standard mass univariate analysis was performed in SPM5 whereby a general linear model (GLM) estimated the response to each sentence in each run. These effects were modeled with a boxcar function convolved with the canonical Hemodynamic Response Function (HRF). The model also included first-order temporal derivatives of these effects (which were not used in the analyses), as well as nuisance regressors representing entire experimental runs and offline-estimated motion parameters.

#### Functional localization

Data analyses were performed on fMRI BOLD signals extracted from the bilateral fronto-temporal language network. This network was defined functionally in each participant using a well-validated language localizer task (Fedorenko et al., 2010), where participants read sentences vs. lists of nonwords. This contrast targets brain areas that support ‘high-level’ linguistic processing, past the perceptual (auditory/visual) analysis. Brain regions that this localizer identifies are robust to modality of presentation (Fedorenko et al., 2010; Scott et al., 2017; Malik-Moraleda, Ayyash et al., 2022), as well as materials and task (e.g., Diachek, Blank, Siegelman et al., 2020). Further, these regions have been shown to exhibit strong sensitivity to both lexico-semantic processing (understanding individual word meanings) and combinatorial, syntactic/semantic processing (putting words together into phrases and sentences) (Bautista & Wilson, 2016; I. Blank et al., 2016; I. A. Blank & Fedorenko, 2020; Fedorenko et al., 2010, 2012, 2016, 2020). Following prior work, we used group-constrained, participant-specific functional localization (Fedorenko et al., 2010). Namely, individual activation maps for the target contrast (here, *sentences>nonwords*) were combined with “constraints” in the form of spatial ‘masks’— corresponding to broad areas within which most participants in a large, independent sample show activation for the same contrast. The masks, which are derived in a data-driven way from this independent sample of participants and are available from https://evlab.mit.edu/funcloc/, have been used in many prior (e.g., Diachek, Blank, Siegelman et al., 2020; Jouravlev et al., 2019; Shain, Blank et al., 2020). They include six regions in each hemisphere: three in the frontal cortex (two in the inferior frontal gyrus, including its orbital portion: IFGorb, IFG; and one in the middle frontal gyrus: MFG), two in the anterior and posterior temporal cortex (AntTemp and PostTemp), and one in the angular gyrus (AngG). Within each mask, we selected 10% of most localizer-responsive voxels (voxels with the highest *t*-value for the localizer contrast) following the standard approach in prior work. This approach allows to pool data from the same functional regions across participants even when these regions do not align well spatially in the common space.

We constructed a stimulus-response matrix for each of the two experiments by i) averaging the BOLD responses to each sentence in each experiment across the three repetitions, resulting in 1 data point per sentence per language-responsive voxel of each participant, selected as described above (13,553 voxels total across the unique 10 participants; 1,355 average, ±6 std. dev.), and ii) concatenating all sentences (384 in Experiment 2 and 243 in Experiment 3), yielding a 384x12,195 matrix for the 9 unique participants in Experiment 2, and a 243x8,121 matrix for the 6 unique participants in Experiment 3.

### ANN models

As our computational models, we chose to investigate the GPT2 Transformer model family (Radford et al., 2019). These models are trained to predict the next token in a large dataset emphasizing document quality (WebText). We focus on this model family for two reasons:

i) as a unidirectional-attention model, GPT2 arguably processes input in a more human-like manner than bidirectional-attention models such as BERT (Devlin et al., 2018), which have access to the yet unseen input in the future context, and ii) previous work has shown that GPT2 in particular seems to accurately capture human brain activity in the language system during the processing of the same linguistic stimuli (e.g., Schrimpf et al., 2021; Caucheteux & King, 2022; Goldstein et al., 2022). We report results for GPT2-xl, the top-performing ANN language model in previous work (Schrimpf et al., 2021) and validate that the findings hold across the GPT2 model family (**Figure SI 1A**) to ensure the robustness of our results to idiosyncratic model features. Hence, our primary ANN language model of interest was *GPT2-xl* (number of layers L=48, hidden size H=1600). Additionally, we tested *GPT*2 (L=12, H=768) and *Distil-GPT2,* a distilled version (Sanh et al., 2019; L=6 H=768). For all three GPT2 models, we used the pretrained models available via the HuggingFace library (Wolf et al., 2020).

### Retrieving ANN model representations

To retrieve ANN model representations, we treated each ANN model (see <u>Methods; ANN models</u>) as an experimental participant and ran similar experiments on them as the one that was run on humans. We retrieved ANN representations for each sentence for each ANN layer (i.e., at the end of each Transformer block). Given that human participants were exposed to the full sentence at once, we similarly computed a sequence summary representation for each sentence. Our primary approach for obtaining a sequence summary representation was using the *last-token representation:* we obtained the representation of the last sentence token (which was always the representation of the final period token “.”) as a sequence summary, given that unidirectional models already aggregate representations of the preceding context (i.e., earlier tokens in the sentence) (see **Figure SI 1B** for generalization to average-token representations of sentences).

To retrieve ANN model representations, we fed sentences to the model sequentially (i.e., sentence by sentence). For the contextualized representations (see <u>Methods; Manipulations of</u> <u>computational experimental design</u>), we grouped sentences by passage to mimic the experimental procedure for human participants and fed the passage context (if any) before but not after each sentence to the ANN model. For the decontextualized representations, we did not feed any passage context to the model.

### Comparison of ANN model representations to brain measurements

Because we were interested in which aspects of the stimulus contribute to high brain predictivity, we compared ANN model representations of systematically manipulated stimuli (see Methods; Perturbation manipulation conditions below) with brain recordings of humans processing the original (intact) version of the sentences (see Methods; fMRI dataset above).

We treated the ANN language model representation at each layer separately and tested how well it could predict human brain recordings (we treated the two experiments in the Pereira et al., 2018 dataset separately but averaged the results across experiments for all plots). Following Schrimpf et al. (2021), we divided the stimuli (i.e., sentences) into an 80%–20% training–held-out split. For each (participant-specific) voxel, we fitted a linear regression model (ordinary least squares) from the ANN’s representations of the training stimuli to that voxel’s corresponding brain recordings for those stimuli. We applied the regression on model representations of the held-out 20% of stimuli to generate predicted brain responses for those stimuli, and then compared predicted versus actual brain responses for that voxel using the Pearson correlation coefficient. This process was repeated five times, holding out a different 20% of stimuli each time. For each voxel, we then took the mean of the resulting five scores to give us that voxel’s mean predictivity score, computed each participant’s median predictivity score across that participant’s voxels, and computed the median and median absolute deviation (m.a.d.) within-participant error within each perturbation condition manipulation category. We report the results for the best-performing layer of the ANN as well as results across layers, for completeness (**Figure SI 2**).

### Estimation of noise ceiling (quantified as brain-to-brain predictivity)

Due to intrinsic noise in biological measurements, we estimated how well the best possible “average human” model could perform on predicting brain responses in single voxels for held-out “target” participants. In our brain-to-brain predictivity estimation, we included the *n*=5 participants that completed both experiments in the Pereira et al. (2018) dataset to obtain full overlap in the materials across participants. Following Schrimpf et al. (2021), the ceiling value was estimated using a three-step procedure (see **SI Methods** for additional details): We (i) iteratively subsampled the data to predict voxel responses in a given “target” participant from the voxel responses of the remaining “predictor” participants, (ii) extrapolated the procedure to a participant pool of infinitely many participants, and (iii) obtained a final ceiling value by aggregating the estimated voxel-wise predictivity ceilings. Via this procedure, we obtained a ceiling value of 0.32 for the Pereira et al. (2018) dataset.

### Perturbation manipulation conditions

For our baseline *Original* condition, we stripped the sentence stimuli (from Experiments 2 and 3 in Pereira et al., 2018) of all sentence-internal punctuation, except for hyphens and apostrophes, and lower-cased all words. This was done to ensure that conditions are as comparable as possible across manipulation conditions (for instance, it is unclear where sentence-internal punctuation should go when sentence word order is perturbed). For a baseline *control* condition, we created length-controlled random word lists, *RandWordList,* by gathering all words across the dataset into a list and replacing every word in every sentence by a random draw (without replacement). For the critical conditions, we applied a range of controlled manipulations to the stimuli used in the original fMRI experiments reported in Pereira et al. (2018). These manipulations can be grouped into three categories: i) word-order manipulations, designed to understand how degrading word order in various ways affects processing, ii) information-loss manipulations, designed to understand how loss of words from a particular part of speech category affects processing, and iii) semantic-distance manipulations, designed to understand how replacing sentences with sentences that are closer vs. further semantically affects processing. Manipulations were applied once to the full dataset. This perturbed dataset was then fed into the ANN language models sentence-by-sentence and contextualized or decontextualized sentence representations were obtained. (As described in <u>Methods; Retrieving ANN model representations</u>, contextualized representations of perturbed sentences were obtained using the sentence’s passage context which was perturbed in the same way as the sentence of interest.)

#### i) Word-order manipulations

For the word-order manipulations, we investigated ANN-to-brain mapping performance across different sentence-internal word scrambling conditions. For five of the word-order manipulation conditions, we followed the material creation procedure described in (Mollica, Siegelman et al., 2020). Specifically, in four of these conditions, word order was scrambled to different degrees by iteratively and randomly choosing 1, 3, 5 or 7 words from the *Original* sentence stimuli and swapping them with one of their immediate word neighbors, leading to the creation of the *1LocalWordSwap, 3LocalWordSwaps, 5LocalWordSwaps, 7LocalWordSwaps* conditions. To ensure that the desired number of local swaps has in fact been achieved (i.e., within the chosen number of swaps, no swap was undone by another), the pairwise edit distance between the original sentence and the scrambled condition was calculated. As reported in Mollica, Siegelman et al. (2020), these local swap manipulations, even for the 7-swap case (*7LocalWordSwaps*), typically preserve local semantic dependency structure, as can be measured by pointwise mutual information (PMI) among nearby words (as detailed below). The fifth (and last) condition which also followed the creation procedure described in Mollica, Siegelman et al. (2020) was a condition where the PMI among nearby words is minimized (the *LowPMI* condition). Here, we assigned the content and function words of every sentence to two lists (creating 4 lists overall: even- and odd-numbered content words, and even- and odd-numbered function words, according to their position in the sentence). These lists were then re-concatenated into a string such that all function words intervened between the content words in the two lists, creating maximal linear distance between combinable content words (i.e., words that were adjacent/proximal in the original sentence).

We also created two additional word-order manipulation conditions that were not investigated in Mollica, Siegelman et al. (2020). The first used a different strategy for minimizing local combinability than the one used in the *LowPMI* condition: the *LowPMIRandom* condition. Here, stimuli were created by generating 10 random permutations of the words within each sentence (which we ensured did not include the versions used in the *1LocalWordSwap-7* conditions) and choosing the perturbation with the lowest PMI score (computed as detailed next). Given that the *LowPMI* condition was the only condition from the original paper that was generated in a deterministic way, the *LowPMIRandom* condition was included to ensure that the models could not exploit the latent generation procedure. The second was a *ReverseOrder* condition, in which the order of the words in the sentence was reversed. This condition ensures maximal linear distance between words in the original and the manipulated string, while preserving the PMI profile of the original stimulus.

A string’s PMI score was calculated using the procedure described in Mollica, Siegelman et al. (2020): For each string, we used a sliding four-word window to extract local word pairs (this is equivalent to collecting the bigrams, 1-skip-grams, and 2-skip-grams from each string). For each word pair, we then calculated its positive PMI score.^1^ Probabilities were estimated using the Google N-gram corpus (Michel et al., 2011) and ZS Python library (Smith, 2014) with Laplace smoothing (α = 0.1). The string’s PMI score was finally calculated by averaging across the positive PMI values for all word pairs occurring within a four-word sliding window (see Equation *1*). The PMI scores for all conditions can be found in **Figure SI 3**.

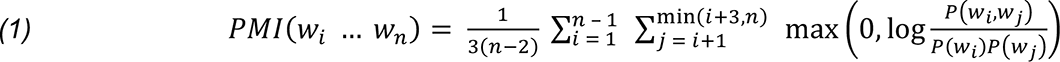

#### ii) Information-loss manipulations

For the information-loss manipulations, we investigated ANN-to-brain mapping performance across five versions of each sentence, for which different subsets of words were retained relative to the original sentence. For the different manipulations, we respectively retained only words whose part of speech tag, as determined by the NLTK part-of-speech tagger (Bird et al., 2009), is in a given set, while preserving the original order of the retained words. Specifically, we examined versions made up of (i) all the content words, i.e., nouns, verbs, adjectives, and adverbs (*KeepContentW)*, (ii) nouns, verbs, and adjectives (*KeepNVA*), (iii) nouns and verbs (*KeepNV*), (iv) nouns (*KeepN*), and (v) only the function words (*KeepFunctionW*). Following O’Connor & Andreas (2021) we included pronouns and proper names in the set of nouns. Note also that because not all the sentences had adverbs and/or adjectives, some pairs of the conditions (i), (ii), (iii), and (iv) could be identical for some sentences.

#### iii) Semantic-distance manipulations

For the semantic-distance manipulations, we investigated ANN-to-brain mapping performance across four conditions, for which the original sentence was replaced by a sentence of variable semantic distances. For three of these conditions, we leveraged the hierarchical organization of the materials in Pereira et al. (2018) (**Figure SI 4**). For the fourth condition, we generated sentence paraphrases. We first describe the three conditions that leverage the hierarchical structure of the Pereira et al. (2018) dataset. As described in <u>Methods; fMRI dataset</u>, stimuli for Experiment 2 consisted of 384 sentences grouped into 96 passages with four sentences each, and stimuli for Experiment 3 consisted of 243 sentences grouped into 72 passages with three or four sentences each. Further, the passages in both experiments came from a smaller number of ‘topics’ that spanned a broad and diverse range of content areas, e.g. *clothes* or *animals*. The 96 passages in Experiment 2 were grouped into 24 topics (with 4 passages per topic; e.g., for the topic of *clothes*, there was a passage about a *dress* and a passage about a *glove*), and the 72 passages in Experiment 3 were also grouped into 24 topics (with 3 passages per topic, e.g., *beekeeping* was a topic, with 3 different passages), non-overlapping with the topics in Experiment 2 (for example passages from each experiment, see **Table SI 1**). We created three experimental conditions. In two of them, *RandSentFromPassage* and *RandSentFromTopic*, each sentence was replaced by a sentence from the same passage or topic, respectively. And in the third, *RandSent*, condition, each sentence was replaced by a sentence that was randomly drawn from the entire dataset, with the constraint that no sentence ended up in its original position (proportion of sentences in *RandSent* condition that come from a different topic than the original sentence: 97.1%; proportion of sentences in *RandSent* condition that come from a different passage than the original sentence: 99.2%).

As described in <u>Methods; Comparison of ANN model representations to brain measurements</u>, the cross-validation scheme used in this paper was a 5-fold cross-validation, holding out 20% of stimuli in each fold. In the *TrainIntact-TestPerturbed* experimental design, the mapping model was trained on ANN representations of stimuli from the *Original* benchmark. When benchmarks by design shuffled stimuli relative to the fMRI data (all semantic-distance benchmarks except *Paraphrase*), this procedure could lead to non-independence in train and test splits. To prevent such overlap between the training and test stimuli in the *TrainIntact-TestPerturbed* versions of these benchmarks, we proceeded as follows: For each of the five cross-validation splits, we i) retrieved the representations of the stimuli that belonged to the test set for the same split in the *Original* benchmark. We then either a) randomly shuffled the order of these activations relative to the fMRI data and ensured that no sentence representation remained in its original position (*RandSent*) or we b) iterated over the passages/topics and, whenever possible (i.e., whenever the test set contained more than one sentence from the given passage/topic), randomly shuffled the sentence representations within the passages/sentences, ensuring that no sentence representation remained in its original position (*RandSentFromPassage/RandSentFromTopic)*. Given this constraint, and because the average number of sentences per passage (4 sentences/passage in Experiment 2, and 3.38 sentences/passage in Experiment 3) was lower than the number of cross-validation splits (n=5), the average percentage of sentences whose representations were not shuffled relative to the associated fMRI data using the default 5-fold cross-validation scheme was 53.72% for *RandSentFromPassage* and 8.97% for *RandSentFromTopic.* Although this procedure led to a high proportion of non-shuffled sentence representations relative to the associated fMRI data in the test set, we opted for this method to ensure consistency and comparability across all *TrainIntact-TestPerturbed* benchmarks, which were thus all trained on the exact same intact sentence representations. To alleviate concerns about the mapping performance being driven mainly by matched fMRI representations, we additionally ran the *RandSent, RandSentFromPassage and RandSentFromTopic TrainIntact- TestPerturbed* benchmark versions, as well as the *Original* and *RandWordList* benchmarks for comparison, using only 2 cross-validation splits instead of the default number of 5 folds. Using this procedure, all but 17.17% of sentence representations could be shuffled relative to its associated fMRI data for *RandSentFromPassage* and all sentences could be successfully shuffled with the associated fMRI data for *RandSentFromTopic,* and the key result pattern was not affected (**Figure SI 5**).

For the fourth and last condition, we generated a paraphrase for each sentence in the set. To do so, we used the online OpenAI ChatGPT interface to generate three paraphrases for each sentence. For each of these paraphrases, we automatically selected the paraphrase that was closest in number of words to the original sentence. These approximately length-matched paraphrases were then manually edited if (i) the absolute difference in number of words between the paraphrased sentence and the original sentence was more than three words, (ii) the paraphrased sentence did not capture the semantic content of the original sentence (as judged by the authors), or (iii) the paraphrased sentence contained different pronouns compared to the original sentence due to ChatGPT history. Out of 627 paraphrase sentences, 111 sentences (17.7%) were manually edited to yield the final set of paraphrased stimuli. Identical to the remaining benchmarks, we stripped the sentence stimuli of all sentence-internal punctuation, except for hyphens and apostrophes, and lower-cased all words. On average, the paraphrased sentences were -0.44 words shorter than the original sentences (median: 0). The paraphrased sentences overlapped partly with the original sentences in terms of their lexical content: the average fraction of overlapping words between the paraphrased and original sentences was 0.46 (median: 0.46, min: 0.05, max: 1).

### Manipulations of computational experimental design

The computational experimental design conditions aim to investigate factors related to how the comparisons between ANN representations and brain data are performed. Specifically, we investigated two factors:

#### i) ANN-to-brain mapping model training stimuli

This condition investigated the effect of the training data for the ANN-to-brain mapping model. In the *TrainIntact-TestPerturbed* condition we trained the mapping model on intact (i.e., original, same as the humans were exposed to) stimuli, and tested the mapping model on perturbed stimuli. In the *TrainPerturbed-TestPerturbed* condition we trained the mapping model on perturbed stimuli, and tested the mapping model on perturbed stimuli (using the same perturbation manipulation type).

#### ii) Contextualization of the linguistic stimuli

This condition investigated the effect of preceding linguistic context on the ANN representations derived for each stimulus according to the structure of the materials investigated (see <u>Methods;</u> <u>fMRI dataset</u>). In brief, humans were presented sentences (one at a time) as part of short (3-4 sentence-long) passages. In the *contextualized* condition, the ANN representations were obtained using the preceding sentences in the passage of interest as context (if any; i.e., the first sentence in a passage would have no preceding contextual information). Because the perturbations were applied to the full set of materials once, and ANN representations were derived based on the perturbed sentences, the preceding context of a sentence was perturbed in the same manner as the sentence of interest^2^. In the *decontextualized* condition, the ANN representations were obtained without any preceding context, and representations were hence obtained using individual, decontextualized sentence representations.

These two factors were crossed in a 2x2 design to yield the four conditions: *TrainIntact- TestPerturbed_Contextualized, TrainPerturbed-TestPerturbed_Contextualized, TrainIntact- TestPerturbed_Decontextualized,* and *TrainPerturbed-TestPerturbed_Decontextualized*.

### Statistical tests

For statistical testing of brain predictivity scores within perturbation manipulation conditions (**Figure 2**, **Figure 4**, **Figure SI 4, Figure SI 5, Figure SI 6**; <u>Results:</u> Section 3.1, Section 3.3), we performed pairwise, two-sided, dependent-samples t-tests for all comparisons among the participant-wise brain predictivity values (i.e., 10 values given that the Pereira et al., 2018 consisted of 10 unique participants) between pairs of conditions. P-values were corrected for multiple comparisons (within each perturbation manipulation condition) using the Bonferroni procedure (i.e., if a perturbation manipulation consisted of 7 conditions and, correspondingly, 7 pairwise comparisons were performed, with each condition compared to the original condition (or to the baseline, random word list, condition) the correction was performed over these 7 tests; for completeness, all pairwise condition comparisons are reported in **Table SI 2**).

**Figure 2.**
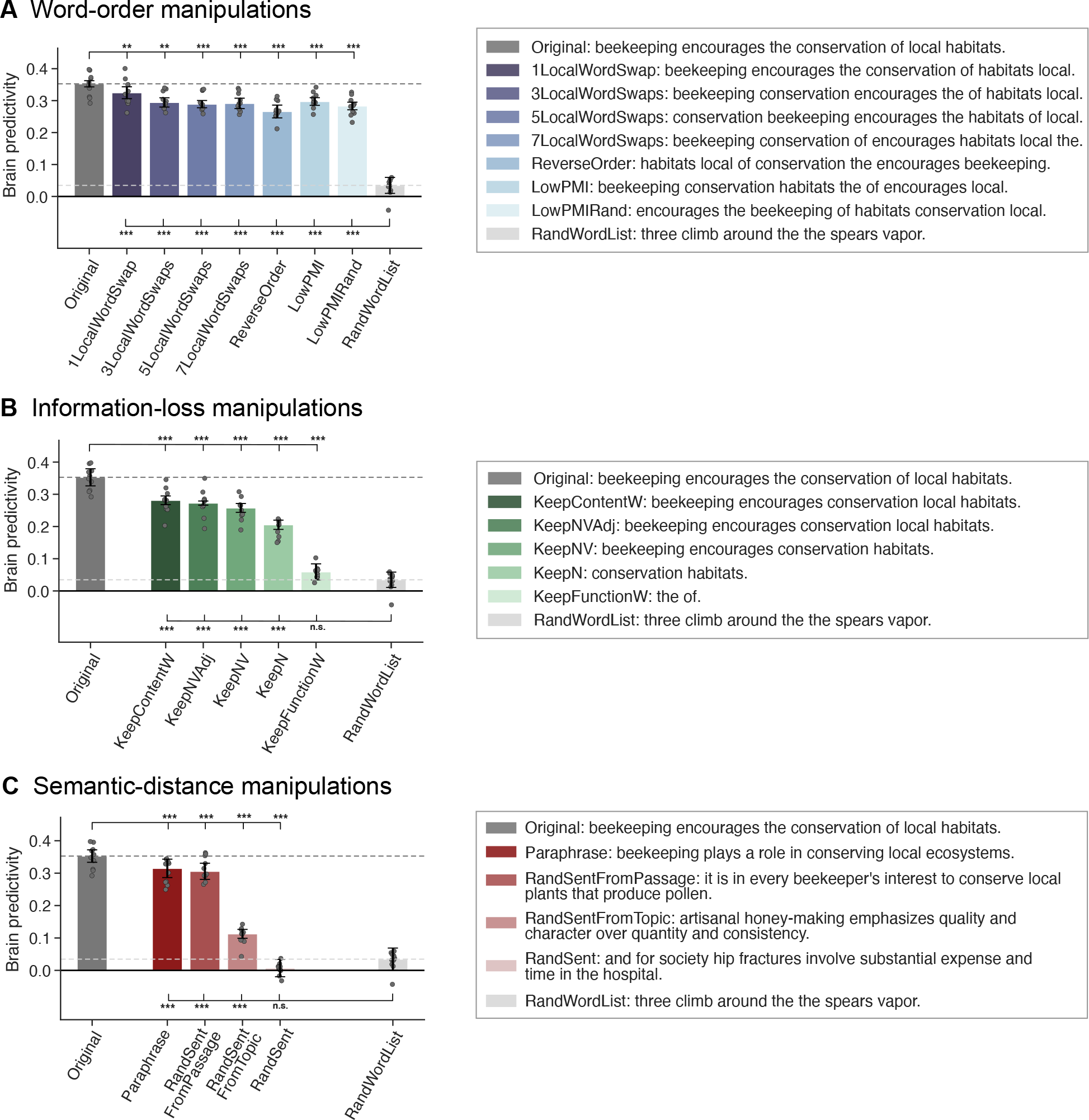
Perturbation manipulations that lead to the loss of lexical-semantic or topical content information decrease brain predictivity. Performance of ANN-to-brain mapping models on held-out sentences, trained on ANN representations of intact sentences and evaluated on ANN representations of perturbed sentences (Figure 1B) from the three perturbation manipulation conditions (panels A-C). For each condition (bar), we plot the raw brain predictivity Pearson r value of the best-performing layer (as in Schrimpf et al., 2021). The ceiling level for the Pereira2018 dataset is r=0.32, as estimated via brain-to-brain predictivity (see <u>Methods; Estimation of ceiling</u>), which for the *Original* benchmark leads to ceiling-level predictivity, in line with Schrimpf et al. (2021). Error bars show median absolute deviation within participants. Manipulation condition scores that were significantly different from the *Original* and *RandWordList* control benchmarks (dark and light gray dashed lines, respectively; these conditions are identical across the three panels) are marked with asterisks (p<.05: *; p<.01: **; p<.001: ***) above and below the bars, respectively, in each graph. Significance was established via dependent two-sided t-tests, with p-values corrected for multiple comparisons (within each perturbation manipulation condition and separately for the comparisons to the original vs. the random word-list baseline) using the Bonferroni procedure.

Error bars of brain predictivity scores show median absolute deviation (m.a.d.) within participants using Scipy 1.8.0’s *median_abs_deviation* function with a scaling factor of ∼0.67 (scale=”normal”) for approximate consistency with the standard deviation for normally distributed data. Thus, error bars were computed by centering the data across conditions within a manipulation category per participant to remove within-participant differences and finally computing the m.a.d. over participants. The error bars hence demonstrate the ANN-to-brain mapping model’s prediction variance within participants across conditions rather than uncertainty around the median.

For statistical testing between computational experimental design conditions (**Figure 6**, <u>Results;</u> Section 3.3), we concatenated the participant-wise brain predictivity values within a perturbation manipulation condition (i.e., if a perturbation manipulation consisted of 7 conditions, we concatenated 10*7 = 70 values). Two-sided dependent-samples t-tests were performed between these pairs of computational experimental conditions. P-values were corrected for multiple comparisons (within each computational experimental design condition) using the Bonferroni procedure. Throughout the figures, significance levels are denoted as follows: p<.05: *; p<.01: **; p<.001: ***.

## 3. Results

### 3.1. Lexical-semantic content, not syntactic structure, is the main contributor to ANN-brain similarity in the language network

In this section, we investigate which aspects of a linguistic stimulus contribute to successful ANN-to-brain correspondence of canonically trained ANN-to-brain mapping models. In particular, we trained ANN-to-brain mapping models on ANN representations of intact stimuli (with corresponding brain responses to intact stimuli) and tested these models using ANN representations of perturbed stimuli (with corresponding brain responses for intact stimuli) (the *TrainIntact-TestPerturbed* approach, as illustrated in **Figure 1B**). We report results for GPT2-xl, the top-performing ANN language model in previous work (Schrimpf et al., 2021) (see **Figure SI 1A** for the generalization of the findings across the GPT2 model family, and **Figure SI 1B** for generalization to a different sequence summarization approach when extracting ANN model representations). In line with Schrimpf et al. (2021), we treat each GPT2-xl layer as an individual model (“layer model”) and report the brain predictivity score for the best-performing GPT2-xl layer model per perturbation condition. We note that the results derived in this way were comparable to selecting the best-performing GPT2-xl model layer on the *Original* benchmark and using this layer for evaluating the remaining perturbation manipulation conditions (**Figure SI 5**). We diverge from Schrimpf et al. (2021) in that the brain predictivity scores throughout the manuscript are *raw* Pearson r values, rather than r values normalized by the noise ceiling value quantified to be r=0.32 via extrapolated brain-to-brain predictivity for the Pereira et al. (2018) dataset (see <u>Methods;</u> <u>Estimation of ceiling</u>).

First, we investigated the performance of the mapping model on a control condition: a length-matched list of random words. For this condition, the mapping model performed at near-chance-level (**Figure 2**, *RandWordList* condition in panels A-C). Chance level (zero predictivity) was not fully reached for the best-performing layer model, possibly because this layer is able to exploit some information about the length of the stimulus (see **Figure SI 6**). We then investigated the effect of our three types of perturbation manipulations—manipulations of word order within the sentence (*Word-order manipulations*), loss of different subsets of words from the sentence (*Information-loss manipulations*), and manipulations of the semantic distance from the original sentence (*Semantic-distance manipulation*)—on the mapping model’s ability to predict brain activity, relative to the original sentence (**Figure 2**, *Original* condition in panels A-C).

#### i) Word-order manipulations (*Figure 2A*)

Word-order manipulations significantly affected brain predictivity, but predictivity scores did not correlate with the severity of word-order manipulations: in particular, predictivity remained relatively high even for the most severe scrambling manipulations, with drops in predictivity values ranging between 8 and 24% for the different manipulations. In particular, one local word swap (*1LocalWordSwap*) led to a ∼8% drop in brain predictivity (0.35 *Original* vs. 0.32 *1LocalWordSwap:* pairwise dependent t-test, t=5.52, p<.01; all reported p-values were corrected for multiple comparisons within each manipulation category using the Bonferroni procedure). The remaining local word swap conditions (*{3,5,7}LocalWordSwaps*) all had a comparable numerical effect (∼17% drop) on brain predictivity (from 0.35 to 0.29, ts>5.81, ps<.001). Pairwise comparisons among the {*1*,*3,5,7}LocalWordSwaps* conditions showed no significant differences (see **Table SI 3** for the pairwise statistical comparisons among all conditions). Even the most extreme local word-order scrambling condition, i.e., reversing the order of the words (*ReverseOrder*), yielded a decrease relative to Original that was statistically comparable to, albeit larger than, the conditions where 3 or more local pairs were swapped (0.35 *Original* vs. 0.27 *ReverseOrder*, ∼24% drop, t=9.29, p<.001).

Critically, all these five conditions (*{1,3,5,7}LocalWordSwaps, ReverseOrder*) were designed to retain local semantic dependency structure as quantified by pointwise mutual information (PMI) (see <u>Methods; Perturbation manipulation conditions; **Figure SI 2**</u>). To test whether preserving local combinability of words is critical for brain predictivity (cf. Mollica, Siegelman et al., 2020), we examined two conditions where local dependency structure was destroyed: the *LowPMI* and *LowPMIRandom* conditions, both of which decreased local PMI. Strikingly, even for these conditions, the effect on brain predictivity was relatively small, similar to the local-scrambling conditions (0.35 *Original* vs. 0.30 *LowPMI*, ∼16% drop, t=6.35 p<.001; and vs. 0.28 *LowPMIRandom*, ∼20% drop, t=8.47, p<.001). Hence, destroying the local dependency structure does not appear to affect brain predictivity beyond how it is affected by local word swaps that keep local dependency structure more easily inferable.

#### ii) Information-loss manipulations (*Figure 2B*)

Preservation of different subsets of content-carrying words relative to the full sentence was associated with relatively high brain predictivity, though all of these conditions led to a significant drop in performance relative to the *Original* condition (0.35 *Original* vs. 0.28-0.21, ts=9.21-28.58, ps<.001; 20-42% drops). Preserving fewer content words—preserving all content words (*KeepContentW*), preserving only the nouns, verbs, and adjectives (*KeepNVA*), only the nouns and verbs (*KeepNV)*, or only the nouns (*KeepN)*—led to a gradual decrease in predictivity values, even though scores for *KeepContentW and KeepNVA* as well as for *KeepNVAdj and KeepNV* did not differ significantly from each other (see **Table SI 3**). By contrast, retaining only the function words (i.e., removing all content-carrying words) led to a brain predictivity comparable to that of a random word list (0.03 *RandWordList* vs. 0.06 *KeepFunctionW*, t=-2.57, p>.05; a ∼83% drop from the *Original* condition). To ensure that the strong drop in predictivity for the *KeepFunctionW* condition was not merely an artifact of the length of the condition (a relatively low number of words in each input string, **Table SI 4**), we included an additional control condition (*RandN*, **Figure SI 7**), which was matched for length with the *KeepN* condition, but in which the nouns were randomly sampled from the nouns in the dataset. This *RandN* control condition was associated with predictivity performance no different than the random word list control condition (0.03 *RandWordList* vs. 0.04 *RandN,* t=-0.96, p>.05), and similar to the *KeepFunctionW* condition (0.04 *RandN* vs. 0.06 *FunctionWords*, t=-1.88, p>.05, **Table SI 2**). These results highlight a large asymmetry in the contribution of content vs. function words to brain predictivity and suggest that preserving more of the lexical semantic content leads to higher predictivity.

#### iii) Semantic-distance manipulations (*Figure 2C*)

As expected, replacing the original sentence with a random sentence was associated with chance-level predictivity (0.01, ∼98% predictivity drop; one-sample t-test to 0: t=0.78, p>.05), similar to that of a random list of words (*RandWordList* vs. *RandSent,* t=2.1, p>.05; ruling out the possibility that any well-formed and meaningful sentence would yield high brain predictivity). Replacing the sentence with a sentence from the same topic was associated with a ∼68% drop relative to *Original* (0.11; *Original* vs *RandSentFromTopic*, t=28.35, p<.001), much lower than the predictivity associated with word order scrambling manipulations (∼8-24% predictivity drop range) or manipulations that preserve at least some of the content words (e.g., ∼42% predictivity drop in the *KeepN* condition). This result demonstrates that a rough topical overlap does not suffice for high brain predictivity. However, replacing the original sentence with a sentence from the same passage was associated with a drop in predictivity of ∼13% (0.31; *Original* vs. *RandSentFromPassage,* t=6.84, p<.001). (Note that in the *RandSentFromPassage* and *RandSentFromTopic* conditions where sentences were shuffled within subparts of the hierarchically structured dataset, an unavoidable overlap between train and test sentences was introduced for the experimental setup using 5 splits. We show that no key pattern of results was affected using a 2-split cross-validation split in **Figure SI 8** and report the results for the 5-fold experimental paradigm here for consistency across manipulation types.) Finally, replacing the original sentence with a paraphrase led to a drop of ∼11% (0.31; *Original* vs. *Paraphrase*, t=10.78, p<.001), which is comparable to the predictivity of the *RandSentFromPassage* condition, even though the lexical overlap is substantially higher between the *Original* and the *Paraphrase* conditions compared to the *Original* vs. the *RandSentFromPassage* condition (**Figure SI 9**). The result from the *Paraphrase* condition shows that a) even sentences that are highly similar in overall meaning are still associated with a small but reliable decrease in predictivity relative to the original sentence, which can be taken to suggest that the model-to-brain match is sensitive to subtle differences in wording, which are associated with subtle semantic differences; and b) when a certain degree of sentence-level semantic similarity with *Original* is reached (as the case for both *Paraphrase* and *RandSentFromPassage* conditions; see **Figure SI 4**), stronger lexical overlap does not have much of an effect on predictivity as evidenced from the fact that *Paraphrase* was not significantly different from *RandSentFromPassage* (**Table SI 3**).

### 3.2. Perturbation manipulations that are associated with larger representational distortion in the ANN embedding space and render linguistic stimuli more surprising lead to lower brain predictivity

In this section, we investigate *why* certain perturbation manipulation conditions yield lower brain predictivity than others. We explore two potential factors: a) differences between the original sentences and the perturbed versions in the ANN representational embedding space (<u>Results;</u> Section 3.2.1), and b) the effect of the perturbation manipulations on the ANN’s task performance (i.e., next-word prediction performance; <u>Results;</u> Section 3.2.2).

#### 3.2.1 Perturbation manipulations that are associated with larger representational distortion in the ANN embedding space lead to lower brain predictivity

Do changes in the ANN representational space across perturbed sentence sets (relative to the intact sentences) explain why certain perturbation conditions yield lower brain predictivity than others? To find out, we investigate—for all ANN model layers—what makes some ANN layer representations more suitable than others for predicting brain responses. In particular, we investigated whether layers for which representations of the perturbed stimuli are more similar to the representations of the intact sentences perform better at predicting brain responses. To do so, for each of 18 perturbation manipulations (1 original, 7 word-order manipulations, 5 information-loss manipulations, 4 semantic-distance manipulations, and 1 control (random word list) manipulation), we calculated the degree of representational similarity (as quantified by the Spearman rank correlation coefficient, ρ) between a layer’s representation of the original, intact sentence and the corresponding perturbed sentence. We then averaged these correlation coefficients across all intact-perturbed pairs, to derive a single value per perturbation manipulation per ANN layer. We then correlated these average correlation values with the associated brain predictivity scores (i.e., a total of 864 values: 18 average correlation values x 48 layers).

We observed a strong positive correlation (Pearson r=0.72 across all perturbation manipulation conditions, p<.001; **Figure 3A**) between i) the similarity of an ANN layer’s representation of the original and perturbed stimuli for a given manipulation and ii) how well that layer could predict neural responses for that perturbation manipulation. The positive relationship differed across perturbation manipulation conditions, but was statistically significant in each condition (**Figure 3A,** panels i-v). This relationship suggests that perturbation manipulation conditions that distort the representation of the original, intact sentences to a larger extent are associated with lower brain predictivity scores. On average, later layers (red colors) yielded higher brain predictivity scores; For some perturbation manipulation conditions, like the *KeepFunctionW* condition (pentagon symbol), however, all layers (including later ones) exhibited poor brain predictivity performance that was also associated with consistently low representational similarity to the intact sentences.

**Figure 3.**
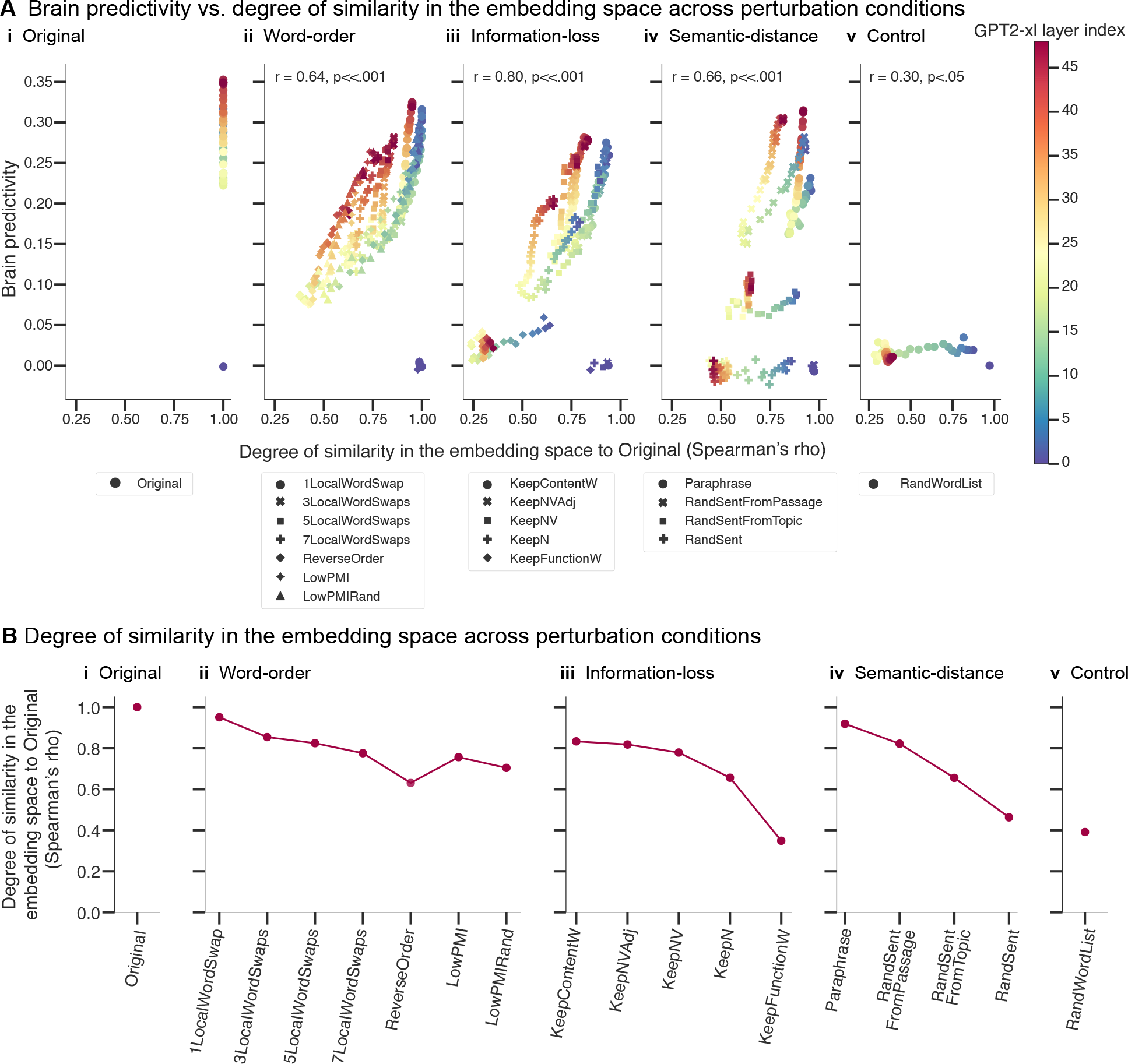
Representational similarity to the original sentences is correlated with brain predictivity. A. Each individual data point shows the correlation between brain predictivity (y-axis) and degree of similarity to the intact sentence set (x-axis, quantified using the Spearman’s rank correlation coefficient, ρ) for a layer of the GPT2-xl ANN model and a certain perturbation manipulation condition. The ANN layer index is denoted by colors. The perturbation manipulation condition is denoted by data point marker symbols. **B.** Similarity of the representations from the last layer of GPT2-xl across conditions to its representations of the intact sentences (note though that the brain predictivity scores reported in the previous sections are from the best-performing layer, not the last one).

**Figure 4.**
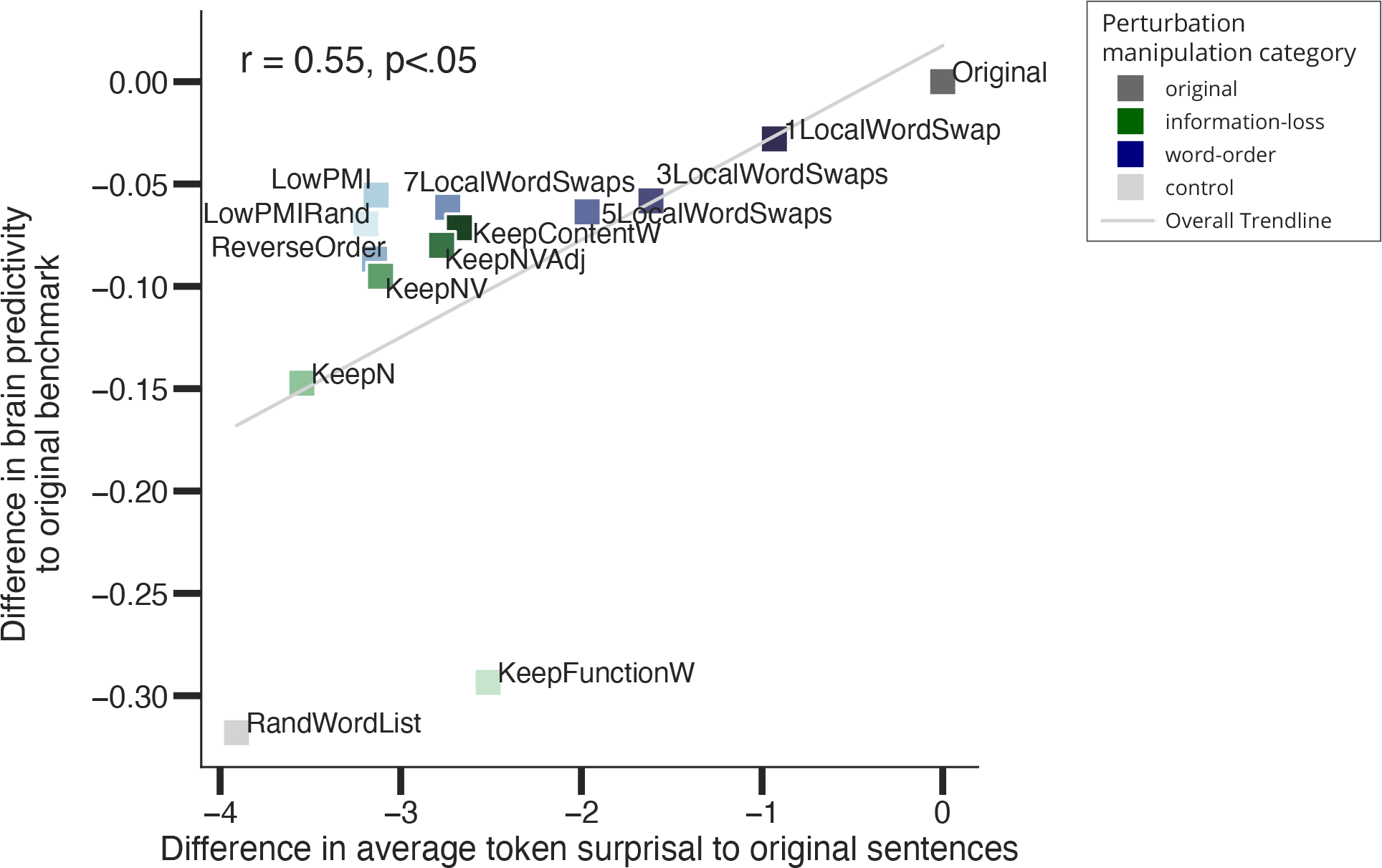
Correlation between difference in brain predictivity relative to the original brain predictivity score and difference in average string surprisal relative to the original sentence condition. Individual data points are perturbation manipulation conditions (original, information-loss, word-order, and control) colored according to overall perturbation manipulation category. Surprisal values are in nats (logarithm to base e). Note that we excluded the semantic-distance manipulation category for this analysis, because 3 out of 4 of these manipulations by design shuffled sentences across the entire material set and hence i) each string did not bear relation to the original string, and ii) average surprisal values across the materials would be identical to *Original*. In contrast, the stimuli in the two other perturbation categories bear relation to the original string: The information-loss conditions retain words of certain parts of speech relative to the intact sentence, word-order manipulations retain all lexical items from the original string, and the length-matched control condition *RandWordList* exchanges every word in the sentence with a different word and is thus length-matched.

Finally, to understand the trends in **Figure 3A** at a finer grain, we investigated the degree of similarity between the representations of the intact and perturbed stimuli in a selected layer (here: GPT2-xl’s last layer) across perturbation manipulation conditions (**Figure 3B**). As expected, relatively subtle manipulations (e.g., *1LocalWordSwap*) did not strongly affect the representational similarity: the representation of the perturbed sentence versions is very similar to that of the original versions (Spearman ρ=0.95). Across the word-order manipulations (**Figure 3B**; panel ii), representational similarity to the intact sentences gradually decreased with the severity of the word-order scrambling. Likewise, across the information-loss manipulations (**Figure 3B**; panel iii), representational similarity decreased the more lexical content was removed, with representations of only the nouns in the sentence already achieving an average representational similarity of 0.66. Across the semantic-distance manipulations (**Figure 3B**; panel iv), representational similarity decreased with increasing semantic distance. The most destructive manipulations (e.g., the *KeepFunctionW, RandSent,* and *RandWordList* conditions) were the least similar in their representations to the original sentences. Note that the random word list control condition, although showing lower similarity to the original sentences than all the critical perturbation conditions (except the *KeepFunctionW* condition), still achieved a similarity score of 0.39. This suggests that GPT2-xl’s representations of length-matched random word lists are not orthogonal to those of intact sentences (i.e., some units in the last layer of GPT2-xl respond similarly independent of the specific words).

The overall pattern across perturbation manipulation conditions, shown in **Figure 3B**, is similar to the pattern of brain predictivity scores shown in **Figure 2**. This similarity mirrors the main finding from **Figure 3A**, which includes information on all perturbations across all ANN layers: perturbation manipulations that render the representations more distinct from those for intact sentences also result in lower brain predictivity scores.

#### 3.2.2 Perturbation manipulations that render linguistic stimuli more surprising lead to lower brain predictivity

In this section, we ask whether the performance of the ANN-to-brain mapping model is linked to the next-word prediction accuracy of a language ANN model. The most widely used training task for large-scale language ANNs is word-in-context prediction, which aims to minimize the surprisal of a word in the input string conditioned on its context. For this analysis, we obtained the average token surprisal of each input string and averaged these surprisal values across items in each linguistic manipulation condition. We then correlated the difference in these average surprisal values for each condition, relative to the surprisal of the original string, with the difference in brain predictivity for each manipulation condition, relative to brain predictivity for the original condition. Sentence surprisal values were always obtained for the last layer of the ANN (given that GPT2 models are trained to predict next tokens using the last layer representation of the context, and not any other layer representation), whereas brain predictivity scores were derived from the best-performing layer, as before.

Across the *Original*, *Word-order, Information-loss, and Control* perturbation manipulation conditions, we observed a positive correlation between the difference in average string surprisal (i.e., surprisal averaged across tokens in a string, and then averaged across items in a condition) and the difference in brain predictivity relative to the *Original* sentence condition (Pearson r=0.58, p<.05), indicating that stimuli with high surprisal yield less predictive ANN representations for encoding brain responses. This finding suggests that sentence perturbations that affect an ANN language model’s mapping onto brain responses also affect the language model’s performance on the next-word prediction task.

### 3.3. The pattern of brain predictivity across linguistic perturbation manipulations is robust to variation in the computational experimental design

In Sections 3.1 and 3.2, we provided a systematic analysis of the aspects of linguistic stimuli that contribute to the high performance of ANN-to-brain mapping models, as reported in Schrimpf et al. (2021) and investigated why certain perturbation manipulations yield lower brain predictivity than others. In this section, we investigate the robustness of these findings to changes in the computational experimental design.

In particular, we focus on two factors of the experimental design: training the mapping model on intact vs. perturbed stimuli and contextualization of sentence representations with respect to the preceding passage context, crossed in 2x2 factorial design (as summarized in **Figure 5**). **Figure 6** (panels A-E) shows each of these four factor combinations as individual, colored lines across our perturbation manipulations. The experimental design condition investigated in <u>Results;</u> Section 3.1 (and <u>3.2</u>) is the *TrainIntact-TestPerturbed_Contextualized* condition (dark purple lines).

**Figure 5.**
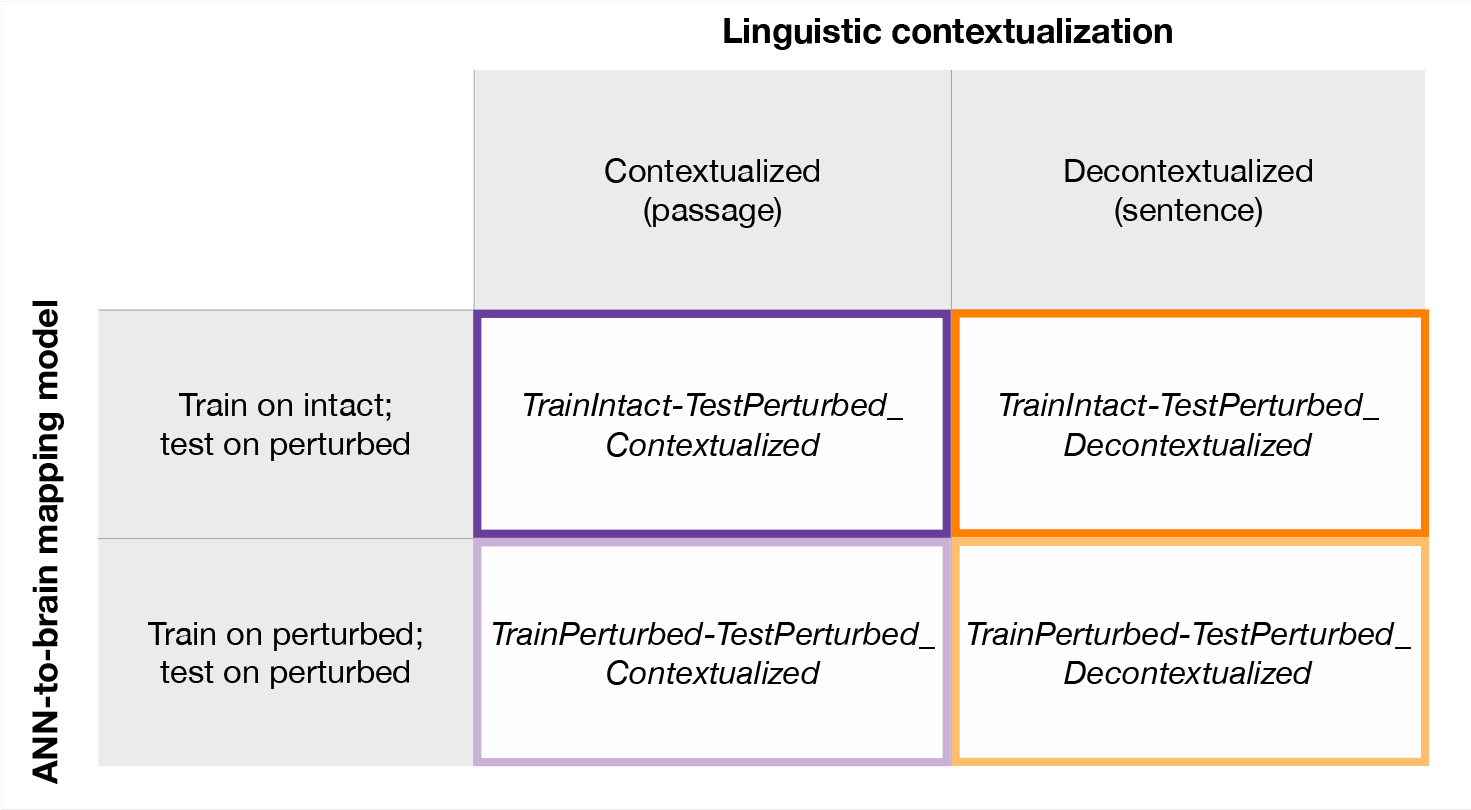
Overview of computational experimental design conditions. Overview of the computational experimental condition factors investigated. We provide a factorial analysis of the contribution of i) the ANN-to-brain mapping model (either forcing the mapping model to generalize to novel perturbation types during test time (*TrainIntact-TestPerturbed*) or allowing the mapping model to exploit perturbation types seen at training time (*TrainPerturbed-Test-Perturbed*), and ii) linguistic contextualization in the computational experimental design (either mimicking the human experimental design and providing prior passage context (*Contextualized*, see <u>Methods; fMRI dataset</u>) or no sentence-external context (*Decontextualized*). We note that the condition presented in <u>Results;</u> Section 3.1.1 and Figure 2 is the *TrainIntact-Test-Perturbed_Contextualized*.

The four computational experimental design conditions yielded highly similar brain predictivity patterns across the 18 perturbation manipulation conditions, as was evidenced by an average pairwise Pearson correlation of r=.84 (p<.001). The lowest pairwise correlation across perturbation manipulations (Pearson r=.63) was obtained by comparing *TrainPerturbed- TestPerturbed_Contextualized* vs. *TrainIntact-TestPerturbed_Decontextualized* while the highest correlation was obtained for *TrainPerturbed-TestPerturbed_Decontextualized* vs. *TrainIntact- TestPerturbed_Decontextualized* (Pearson r=.96).

We note that even though all computational experimental conditions were highly correlated, there was a substantial difference in the magnitude of brain predictivity scores associated with each profile. For example, brain predictivity for the intact (*Original*) condition ranged between 0.26 and 0.35, (**Figure 6A**, *Original*). Across perturbation manipulation conditions, we observed a boost in brain predictivity performance when including previous in-passage sentences as context (purple lines vs. orange lines): on average brain predictivity improved by .06 for *TrainIntact-TestPerturbed* designs and .14 for *TrainPerturbed-TestPerturbed* designs (cf. <u>Discussion;</u> Section 4.3) (see **Table SI 5** for all pairwise comparison statistics between computational experimental design profiles within a manipulation condition).

**Figure 6.**
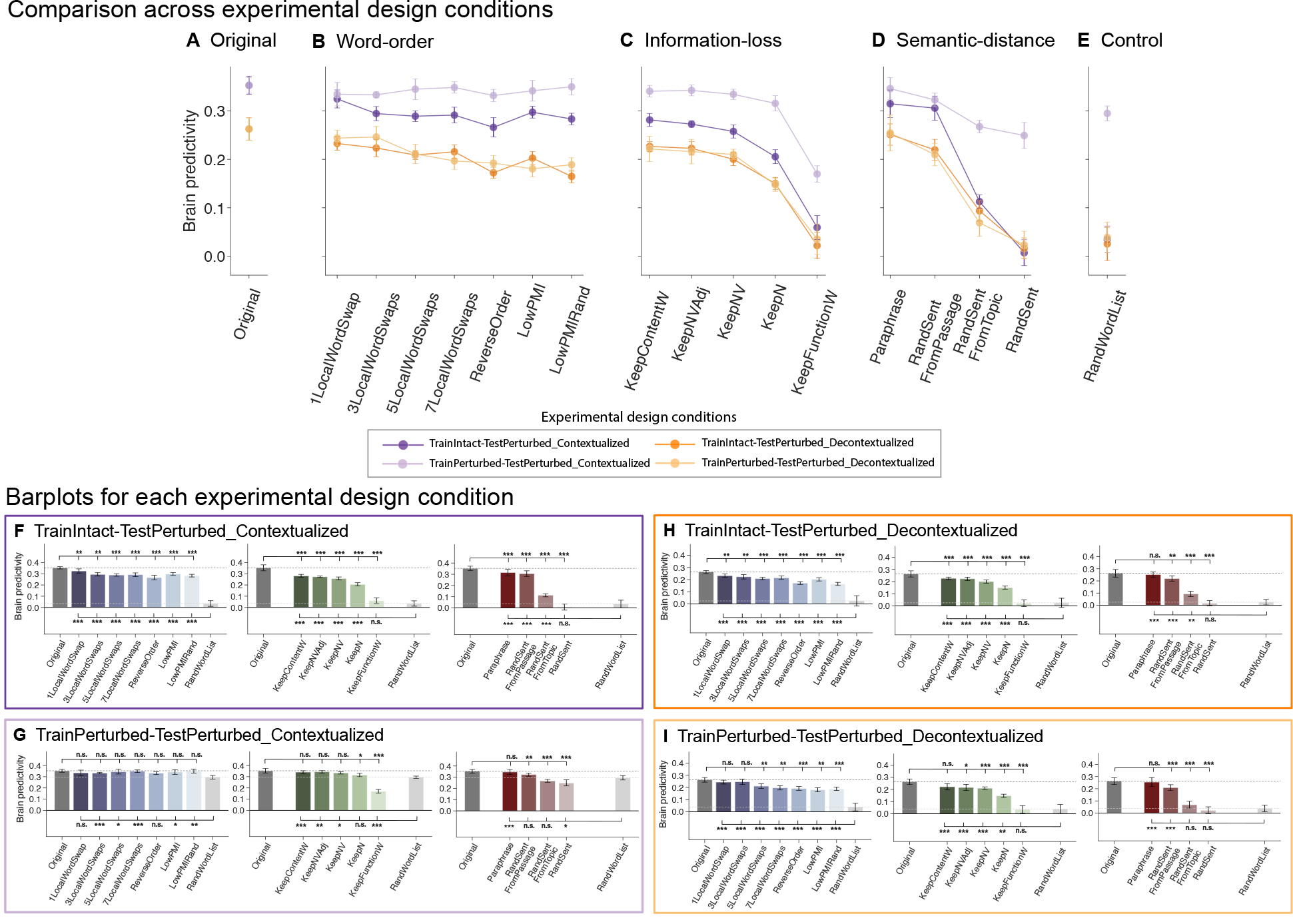
Brain predictivity patterns are largely robust to variations in the computational experimental design. A-E. Comparison of brain predictivity across experimental design conditions (as summarized in Figure 5). Each experimental design condition is shown as an individual line across our perturbational manipulation conditions (individual panels). For each condition, we plot the raw brain predictivity (Pearson r) of the best-performing layer, i.e., the fraction of variance explained that the model can predict relative to the ceiling of the fMRI dataset. We note that the condition presented in <u>Results;</u> Section 3.1 and Figure 2 is the *TrainIntact-TestPerturbed_Contextualized* condition (dark purple line). **F-I**. Barplots for each experimental design condition. Each panel (with a series of barplots) corresponds to a single line in panels A-E with box color matching the line color. Manipulation condition scores that were significantly different from the *Original* and *RandWordList* control benchmarks (dark and light gray dashed lines, respectively) are marked with asterisks (p<.05: *; p<.01: **; p<.001: ***). Significance was established via dependent two-sided t-tests, with p-values corrected for multiple comparisons (within each perturbation manipulation condition) using the Bonferroni procedure. Error bars show median absolute deviation within participants.

For all experimental design conditions, we observed a substantial drop in ANN-to-brain mapping performance for the random word list control condition relative to *Original* (**Figure 6E**, *Control*) (*Original* vs. *RandWordList*: p<.05 across all four factor combinations). Nevertheless, when the mapping model was trained *and* tested on contextualized representations of random word lists, the performance of the mapping model was unexpectedly high (*TrainPerturbed- TestPerturbed_Contextualized*, light purple line). For the same computational experimental design, we also observed surprisingly high ANN-to-brain mapping model performance for the random sentences semantic-distance manipulation (*RandSent*). These results suggest that this mapping model was still able to extract a substantial amount of useful information for predicting responses to held-out sentences, even for stimuli that we would not expect to carry a lot of useful information for match-to-brain (indicating an undesired interaction between the contextualization and cross-validation schemes, see <u>Discussion</u>). However, not just *any* input sentence elicited a high brain predictivity score in this design. Replacing all words in the sentence with one and the same word (**Figure SI 6B**, condition *LengthControl)*, led to a near chance-level performance (see also **Figure SI 4)**. This result shows that, when allowed to exploit meaningful, lexical semantic content from the context, a mapping model can use ANN representations derived from random word lists and random sentences to obtain relatively high predictivity; even though low-level features of the stimulus, such as its length, are not sufficient to obtain high predictivity (**Figure SI 6B**).

In sum, our findings suggest that the conclusions from <u>Results:</u> Section 3.1 regarding the contributions of features of the linguistic input are mostly robust against variation in the computational experimental design. In the Supporting Information, we report results indicating that the conclusions of Section 3.2 are similarly robust: across computational experimental design conditions, we observed that greater linguistic perturbations lead to a) more divergent representations in the ANN’s embedding space (relative to the representations of intact sentences; **Figure SI 10**) and b) a decrease in the ANN’s next-word prediction task performance, i.e., its ability to predict upcoming tokens in those stimuli (**Figure SI 11**).

## 4. Discussion

A number of independent studies have recently shown that representations from state-of-the-art ANN models—especially unidirectional Transformer models—align well with brain responses of humans processing linguistic input (e.g., Jain & Huth, 2018; Pereira et al., 2018; Toneva & Wehbe, 2019; Gauthier & Levy, 2019; Schrimpf et al., 2021; Caucheteux & King, 2022; Goldstein et al., 2022; Kumar et al., 2022; Merlin & Toneva, 2022; Millet et al., 2022; Oota et al., 2022; Pasquiou et al., 2022). However, what makes ANN representations align with human neural responses to language has been little explored (cf. Gauthier & Levy, 2019; Merlin & Toneva, 2022; Oota et al., 2022). Focusing on the top-performing unidirectional-attention GPT model family (Schrimpf et al., 2021), we systematically investigated the effect of diverse linguistic perturbation manipulations, including manipulations that strongly affect sentence meaning (carried largely by content words) and those that primarily affect syntactic structure (carried by word order and function words) on the ability of an ANN-to-brain model to predict brain responses.

The contributions of our work are three-fold: First, we found that lexical semantic content is a stronger contributor to the similarity between ANN language models and brain data than syntactic structure (conveyed by word order or function words), although above a certain level of sentence-level semantic similarity, lexical overlap no longer contributes much. Second, we found that linguistic perturbations that decrease brain predictivity have interpretable causes: they lead to more divergent representations in the ANN’s embedding space (relative to the representations of intact sentences) and decrease the ANN’s next-word prediction task performance, i.e., its ability to predict upcoming tokens in those stimuli. Finally, we found that the effects of these linguistic manipulations are largely robust to variations in the ‘computational experimental design’, including whether the mapping model is trained on intact vs. perturbed stimuli and whether the model is fed contextualized representations that mimic the experimental set-up in human experiments. We elaborate on our findings and discuss their implications below.

### 4.1 Lexical semantic content, not syntactic structure, is the primary driver of the ANN-to-brain similarity

We showed that ANN language models exploit the lexical semantic content of the sentence, rather than the sentence’s syntactic form (conveyed via word order or function words) when predicting brain data. Similarly, sentences elicit higher brain predictivity the more topically related they are to the stimulus for which brain representations were obtained. We demonstrated that this pattern is robust across variations in the computational experimental design, indicating that the ANN-to-brain mapping model pays only limited attention to the part of the ANN representation of the stimulus that is sensitive to syntactic information, but rather relies on the representations of the content words’ meanings. These findings align with two growing bodies of evidence: i) one from (computational) neuroscience that points to the relatively greater importance of meaning for both the magnitude and distributed patterns of activation in the brain’s language system as measured/measurable by fMRI (e.g., Fedorenko et al., 2016; Huth et al., 2016; Pereira et al., 2018; Gauthier & Levy, 2019; Mollica, Siegelman et al., 2020), and ii) another from NLP that shows that ANNs do not necessarily need to use word-order information to solve many current natural language processing benchmark tasks (e.g., Pham et al., 2021; Sinha et al., 2021; O’Connor & Andreas, 2021; cf. Abdou et al., 2022; Lasri et al., 2022).

In our study, we aimed to integrate neuroscientific and NLP perspectives on the role of lexical semantic content vs. syntactic information in the building of linguistic representations, and our perturbation conditions took inspiration from both of these fields. Specifically, the word order manipulations investigated here were inspired by an fMRI study that found the human language network responds as robustly to strings with scrambled word order as to naturalistic input as long as the scrambled order still allows for local composition of words into chunks and phrases (Mollica, Siegelman et al., 2020). Similar to these findings, we found that word-order perturbations that preserve local pointwise mutual information lead to only a small decrease in a model’s ability to predict brain responses; but unlike the results reported for human participants in (Mollica, Siegelman et al., 2020), we found that even extreme word-order perturbations, which disrupt local semantic and syntactic dependencies, lead to a similarly small decrease in ANN-to-brain mapping performance. Further, we found that the omission of function words does little to decrease brain predictivity.

We hypothesize that these results are due to the fact that mapping models do not strongly rely on syntactic information (as argued above). However, an alternative explanation is that—at least in the word-order scrambling manipulations—ANN language models might implicitly (albeit perhaps noisily) reconstruct the original sentence (Malkin et al., 2021; Sinha et al., 2021), plausibly enabled by their extensive memory capacity. In particular, ANNs have access to the exact words in the sentence context (up to a maximal token length, which is not exceeded in our sentence material), whereas memory limitations in humans lead them to discard the exact word sequences after extracting the relevant *meaning* from them (e.g., Potter et al., 1998; Potter, 2012a; Christiansen & Chater, 2016; Hahn et al., 2022).

Of course, the inability of ANN-to-brain mapping models to detect the fine-grained structure of the original sentence could also be due to other reasons, such as the low temporal resolution of fMRI data, which might impose limitations on the detection of structure effects. Given that the language system is strongly sensitive to syntactic processing difficulty (e.g., Blank et al., 2016; Shain, Blank et al., 2020; Shain et al., 2022), it is plausible that modeling linguistic representation construction word-by-word (cf. for the whole sentence at once as we did here), along with perhaps using more temporally-resolved data (e.g., from intracranial human recordings), would reveal stronger effects of syntactic structure than the ones found here. Nevertheless, our results show that syntactic structure is not critical in matching ANN representations with fMRI BOLD responses, at least for the summary representations of sentences.

It is also worth noting that stronger effects of structure might be detected in sentence materials where structure is critical to interpretation, as in cases where word order is the only cue to the propositional meaning, in the absence of animacy/plausibility cues (e.g., *The boy introduced the teacher to the girl*) or in cases where the identity/location of a particular function word is critical, again in the absence of plausibility biases (e.g., *He went out of the building* vs. *He went into the building*; or *The book is on the table* vs. *The book is under the table*).

### 4.2 Perturbations that decrease brain predictivity have interpretable causes

Given that different perturbation conditions affected brain predictivity to quite different extents, we investigated potential reasons for these differences and identified two interpretable correlates, one related to the ANN representational space and the other related to ANN task performance. Perturbation manipulations that led to lower brain scores also (i) led to more divergent representations in the ANN’s embedding space (relative to the representations of intact sentences), and (ii) decreased the ANN’s next-word prediction task performance, i.e., its ability to predict upcoming tokens in those stimuli.

Related to the *ANN representational space*, there has been interest in understanding how the units that make up the representations of current large ANN language models change across stimuli and model layers (Ethayarajh, 2019; Biś et al., 2021). In <u>Results;</u> Section 3.2.1 we quantified the changes in the representational space across our perturbation conditions relative to the intact stimuli and found that perturbations that changed the ANN representation to a greater extent (relative to the representation of the original, intact sentence) also led to larger decreases in brain predictivity scores. Interestingly, even for the most extreme perturbations (e.g., replacing a sentence with a random word list or a random sentence) led to representations that were still moderately correlated with the representations of the intact stimuli (even if these altered representations could not capture human neural responses under most computational experimental design settings). This pattern suggests that in our mapping models, only a subset of the full ANN representational space is being used to represent the stimuli investigated here (naturalistic sentences of length 5-20 words and their perturbed versions). More diverse linguistic materials (e.g., sentences of different length, style, content or context length) may engage a larger subset of the ANN representational space in mapping models.

Related to the *ANN task performance*, there has been interest in understanding how the next-word prediction performance of ANN language models is related to ANN-to-brain correspondence (e.g., Schrimpf et al., 2021; Antonello & Huth, 2022; Caucheteux & King, 2022; Goldstein et al., 2022; Hosseini et al., 2022; Merlin & Toneva, 2022), motivated by substantial evidence for predictive processing in human language comprehension (e.g., Rayner et al., 2006; Demberg & Keller, 2008; Bicknell et al., 2010; Smith & Levy, 2013; Henderson et al., 2016; Willems et al., 2016; Lopopolo et al., 2017; Heilbron et al., 2019; Brothers & Kuperberg, 2021; Heilbron et al., 2022; Shain, Blank et al., 2022). Here, we presented evidence that among our perturbations, those that rendered stimuli less predictable, on average, led to larger decreases in brain predictivity performance (see Merlin & Toneva, 2022 for a similar claim for a naturalistic narratives fMRI dataset). This pattern suggests that less predictable strings may yield representations that have features that are less suitable for predicting fMRI brain data.

### 4.3 How close are we to quantitatively accurate and generalizable models of the human language network?

We identified features of linguistic stimuli (namely, lexical semantic content) that ANN-to-brain mapping models exploit when learning a successful mapping to brain responses. These features are exploited by the mapping model independently of whether the mapping model is trained on intact or perturbed stimuli, and of whether the ANN representations of the target sentences are contextualized with respect to the preceding sentences in a passage. At the same time, we demonstrated that these design choices substantially affect the *magnitude* of the brain scores which may lead to different conclusions about the similarity of current ANN language model representations to the ones in the human brain (**Figure 6**; *Original*). Furthermore, we showed that certain computational experimental designs lead to high brain scores for perturbations that we would not expect to not carry a lot of informative structure such as a random word list (*TrainPerturbed-TestPerturbed, RandWordList;* see **Figures 2, 6**).

The findings from the computational experimental design manipulations yield three important insights. First, until the effort of relating ANN model representations to neural representations reaches maturity—and the field (hopefully) agrees on a unified framework for performing model-to-brain comparisons—any findings about the similarity between ANN and human representations should be evaluated for robustness to the details of how the comparisons are performed. Contextualization of stimuli with respect to the preceding linguistic context may be especially important as it may introduce non-independence issues under certain cross-validation set-ups, as elaborated in the third point below.

Second, these findings highlight the general importance of using careful stimulus-based controls (e.g., replacing the stimuli with random sentences or lists of words) when evaluating ANN-to-brain mappings, in addition to using control (e.g., untrained) models. Only examining ANN-to-brain mapping performance for the original stimuli (those presented to human participants) may lead to flawed inferences about the nature of the similarity. For example, if the ANN representation of a list of random words leads to a similar level of mapping performance with a neural response to some sentence as the representation of that sentence, then we cannot infer that the ANN model is representing the sentence in a similar way to humans.

And third, combining the two previous points, the fact that certain computational experimental designs achieve high predictivity performance on stimuli that are not well-matched with the input to humans showcases an important point that has not received sufficient attention in the recent ANN-to-brain literature: contextualizing sentences by including the preceding sentences in the story/passage, to match how the stimuli were presented to humans can lead to inflated brain predictivity performance under certain cross-validation set-ups. In particular, current language models have the ability to keep track of extended contexts, and if contextualization is not properly controlled for, shared context windows for sentences that go into the train set vs. the test set can lead to ‘leakage’ of statistical regularities in these contextualized ANN representations, leaving the two sets not truly independent. Furthermore, on the brain side, neural responses to coherent texts can be correlated across time for (at least) two reasons: a) the participant is still thinking about the content of the previous context when processing the current word/sentence, and/or b) neural measurements tend to be more similar when they are temporally close (the property known as autocorrelation, which is especially prevalent in methods like fMRI that rely on slow physiological changes; e.g., Bullmore et al., 1996). The two sources of statistical leakage (one in the contextualized sentence representations, one in the neural signal) can be potentially exploited by the ANN-to-brain mapping model.

The passage structure of the benchmark we used in the current study (Pereira et al., 2018) allowed us to perform an exploratory analysis of this issue. Given that contextualization affects sentences and the fMRI BOLD signal *within* passages, but not *across* them, we split the stimuli into train and test sets in two ways: by sentence vs. by passage. By-sentence splitting is the approach that was adopted in Schrimpf et al. (2021) and that we followed here; this approach disregards the passage structure and is therefore subject to the ANN contextualization leakage problem just described. In contrast, by-passage splitting, whereby all the sentences from the same passage end up in the same set (train or test) rather than being split across those sets, should solve the leakage problem (although note that this splitting approach *additionally* requires generalization to new semantic domains: e.g., predicting neural responses to sentences about beekeeping when the mapping model has never seen any sentences related to beekeeping).

We found that splitting the train and test sets by passage yielded much lower brain predictivity scores than splitting the dataset by sentences: ∼0.10 brain predictivity (**Figure SI 12)**, in comparison with ∼0.35 brain predictivity for by-sentence splitting when preceding within-passage sentences are included as context, and ∼0.26 for by-sentence splitting when preceding sentences are not included as context. In addition, representations of random sentences and random lists of words are no longer predictive of human neural responses under this splitting approach in the *TrainPerturbed-TestPerturbed_Contextualized* experimental design (in contrast to the same design, i.e., the light purple datapoints in Figure 6a). As laid out above, this drop in predictivity could be due to the following non-mutually exclusive factors: ANN contextualization leakage, fMRI autocorrelation, and/or the greater difficulty of generalizing to novel semantic domains. Given that non-contextualized sentence representations achieve predictivity of ∼0.26 (substantially higher than ∼0.10) (**Figure 6; Figure SI 12**), we can tentatively rule out the contextual leakage in ANN representations as the main contributor. Understanding the contributions to higher predictivity in the by-sentence cross-validation approach of a) temporal autocorrelation in the fMRI signal vs. b) the relative difficulty of generalizing to new semantic domains may require additional data collection (e.g., neural responses to semantically diverse sentences, similar to Pereira et al. (2018), but presented in a random order instead of in passage structure; removing the autocorrelation between semantically related sentences in this new benchmark would enable a direct comparison of generalization of the mapping model to sentences from the same/similar semantic domains vs. to sentences from new semantic domains, and comparing the results with those from the current benchmark would allow quantifying the contribution of autocorrelation to neural predictivity). Regardless of what these future investigations reveal, however, it seems clear that current ANN language models still have much room for improvement before they can serve as accurate and generalizable models of the fMRI BOLD responses in the human language network.

## 5. Conclusion

In this work, we asked why representations from state-of-the-art ANN language models align with human brain responses (as measured with fMRI) during language processing. To do so, we performed a systematic, large-scale investigation of which linguistic features (across three manipulation categories and four computational experimental designs) reliably contribute to ANN-to-brain mapping performance. We found that the ANN-to-brain mapping model mainly attends to the lexical semantic content—the key contributor to the sentence’s *meaning*—rather than to word order or function words, which jointly create the sentence’s *syntactic frame*. Changes in lexical semantic content, compared to word order or function words, lead to more divergent representations in the ANN’s embedding space and also decrease the ANN’s ability to predict upcoming tokens in those stimuli. This pattern of results is robust to variations in the computational experimental design, suggesting that the lexical semantic content of a sentence is reliably encoded in fMRI responses to language. However, our exploratory investigation of different cross-validation settings has also revealed that although current ANN-to-brain mapping models capture a non-trivial amount of variance in human neural data, they do not easily generalize to new semantic contexts, which leaves room for future work to make language models more human-like.

## Acknowledgments

We thank Martin Schrimpf for his time and help with the Brain-Score framework, and Joe O’Connor for helpful discussions. CK gratefully acknowledges support from the K. Lisa Yang Integrative Computational Neuroscience (ICoN) Center at MIT. GT gratefully acknowledges support from the Amazon Fellowship from the Science Hub (administered by the MIT Schwarzman College of Computing) and the International Doctoral Fellowship from American Association of University Women (AAUW). RPL gratefully acknowledges support from a Paul and Lilah Newton Brain Science award and NSF award BCS-2121074. JA gratefully acknowledges support from the MIT-IBM Watson AI Lab, a Sony Faculty Innovation Award, an Amazon Research Award. EF gratefully acknowledges support from NIH awards R01-DC016607, R01- DC016950 and U01-NS121471, as well as research funds from the McGovern Institute for Brain Research, the Brain and Cognitive Sciences department, the Simons Center for the Social Brain, and the Middleton Professorship. RPL, JA, and EF additionally acknowledge support from the MIT Quest for Intelligence Initiative.

## Supplementary Information (SI)

### Supplementary Figures

**Figure SI 1.**
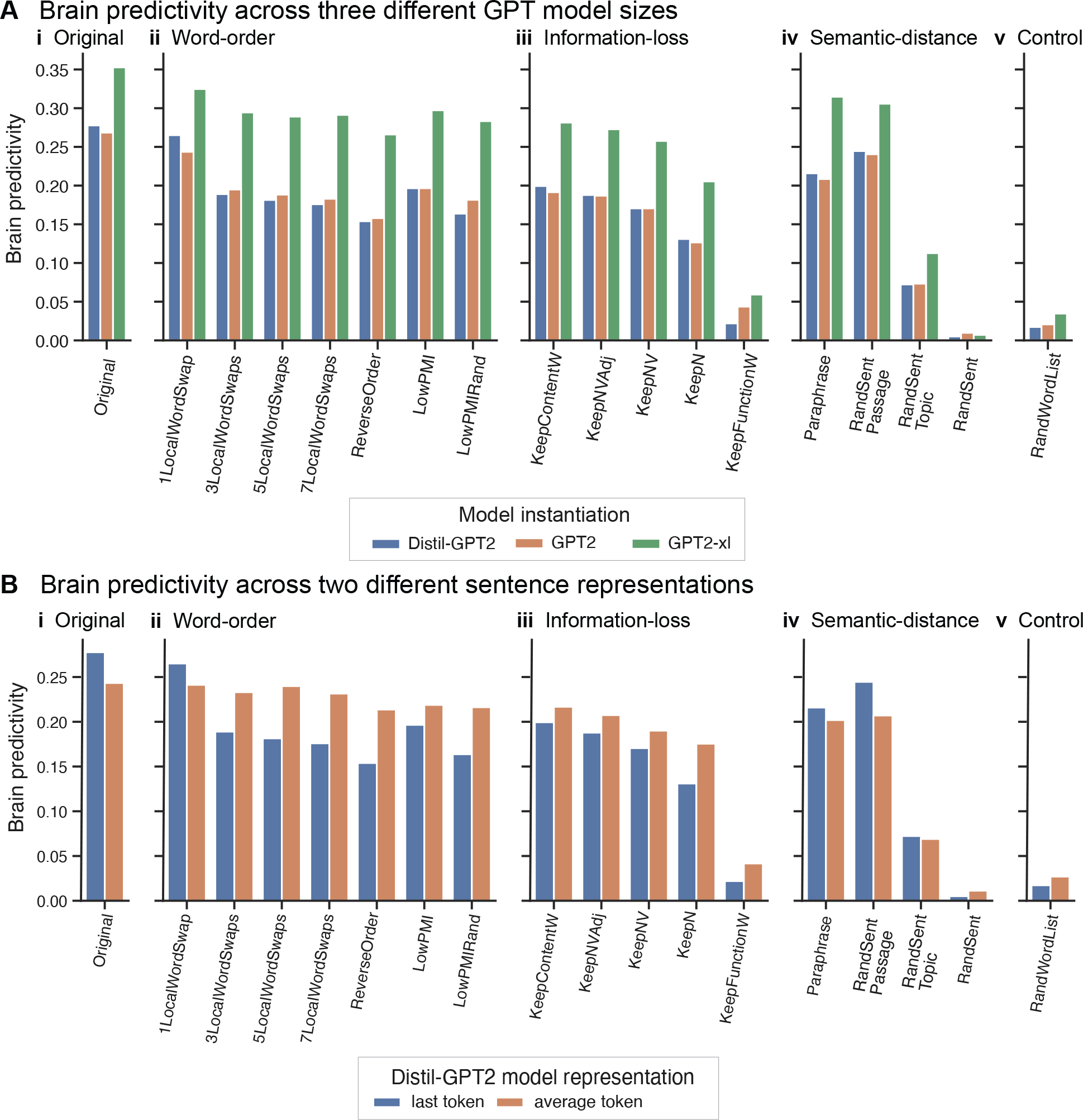
Robustness of brain predictivity scores across ANN model sizes and sentence representations. Across the three model instantiations and the two sequence summary versions, we observe a robust result pattern, with consistent numerical differences between models across conditions. **A.** Predictivity performance of ANN-to-brain models trained on ANN representations of intact sentences and tested on ANN representations of perturbed sentences for three GPT-2 model instantiations (*TrainIntact-TestPerturbed_Contextualized)*: Distil-GPT2, GPT2, and GPT2-xl. **B.** Predictivity performance using two different approaches for representing the sentence for Distil-GPT2 (analyses performed using Distil-GPT2 due to computational cost and robust patterns across ANN model sizes, see panel A). The primary approach was the last token representation where the sequence representation is obtained at the last sentence token (see <u>Methods; Retrieving</u> <u>ANN model representations</u>). We investigated an alternative approach, the average token representation, where we computed the arithmetic mean of all the token representations in the sentence, excluding the token representations of the preceding sentences (if any): in particular, if a sentence was the second sentence within a passage, we did not take into account the token representations from the first sentence in that passage, only the tokens that comprise the current sentence.

**Figure SI 2.**
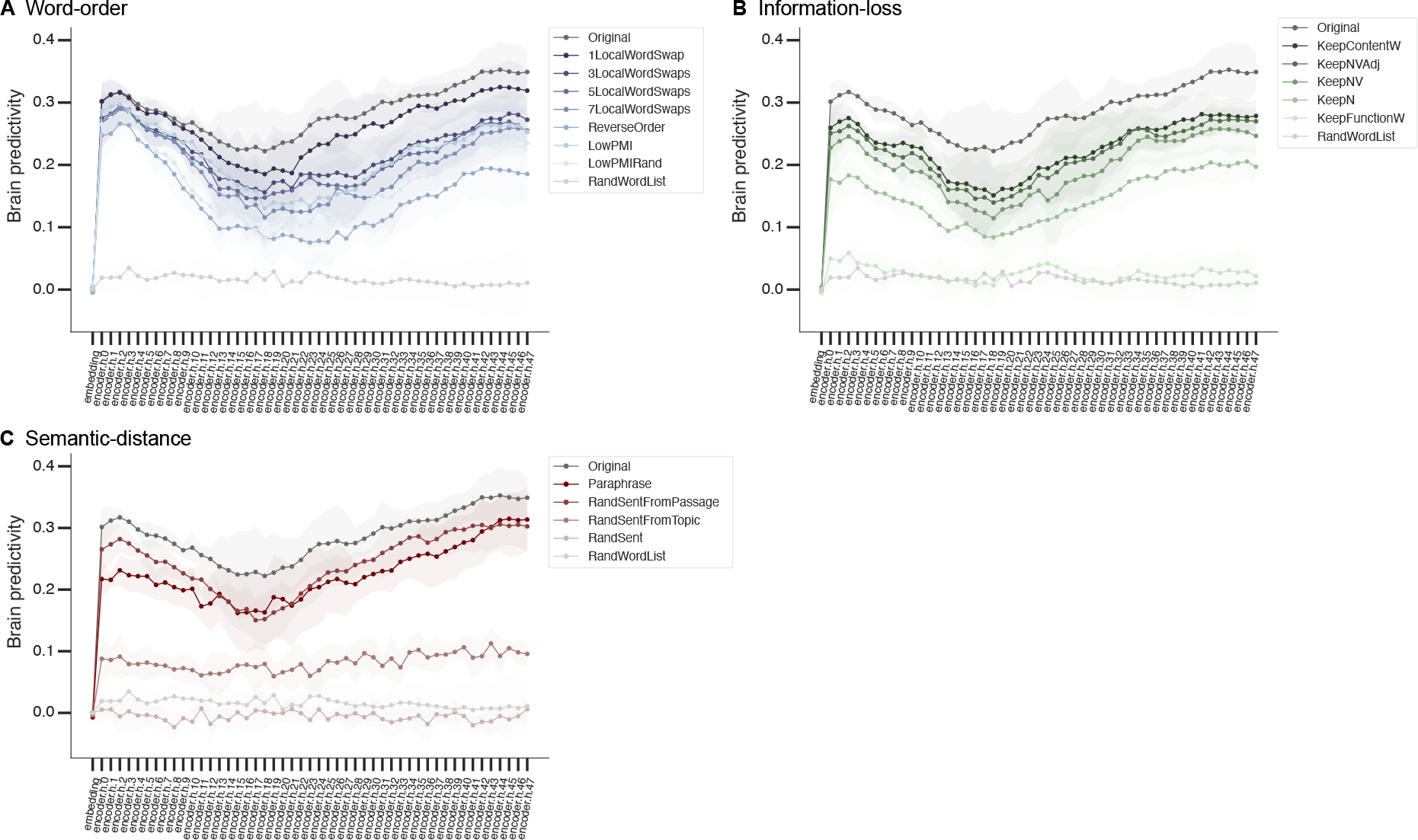
Brain predictivity scores across GPT2-xl layers for each perturbation manipulation condition. Each panel (A-C) shows the scores across layers for all perturbation manipulation conditions within each category. The mapping model was trained on ANN representations of intact sentences and evaluated on ANN representations of perturbed sentences (*TrainIntact-TestPerturbed_Contextualized*). The shaded regions illustrate the median absolute deviation (m.a.d.) error within participants.

**Figure SI 3.**
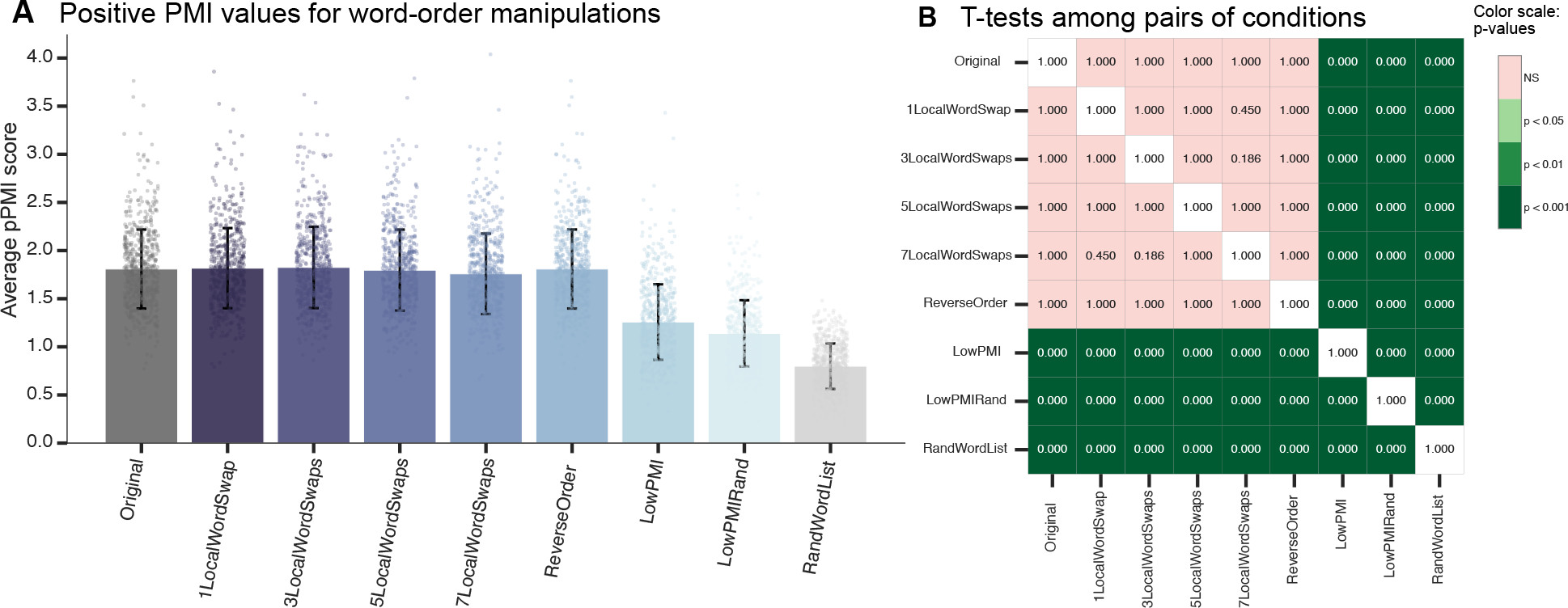
**PMI verification of word-order manipulation conditions**. **A.** Average positive pointwise mutual information (PMI) for each word-order manipulation condition. Each data point represents a sentence, and the error bars show the standard deviation from the mean. The {*1,3,5,7}LocalWordSwaps* and *ReverseOrder* conditions were designed to preserve local dependency structure (see <u>Methods; Perturbation</u> <u>manipulation conditions; Word-order manipulations</u>). As expected, the *Original, {1,3,5,7}LocalWordSwaps* and *ReverseOrder* conditions were not significantly different from each other (panel B). The two low-PMI conditions (*LowPMI* and *LowPMIRandom*) were designed to destroy local dependency structure. As expected, the deterministically created low-PMI condition (*LowPMI*), the nondeterministically-created low-PMI condition (*LowPMIRandom*) and the random wordlist condition (*RandWordList*) were each significantly different from all other conditions. **B.** Significance was established via independent two-sided t-tests, with p-values corrected for multiple comparisons (within each perturbation manipulation condition) using the Bonferroni procedure, here shown as a grid of pairwise p-values for all comparisons.

**Figure SI 4.**
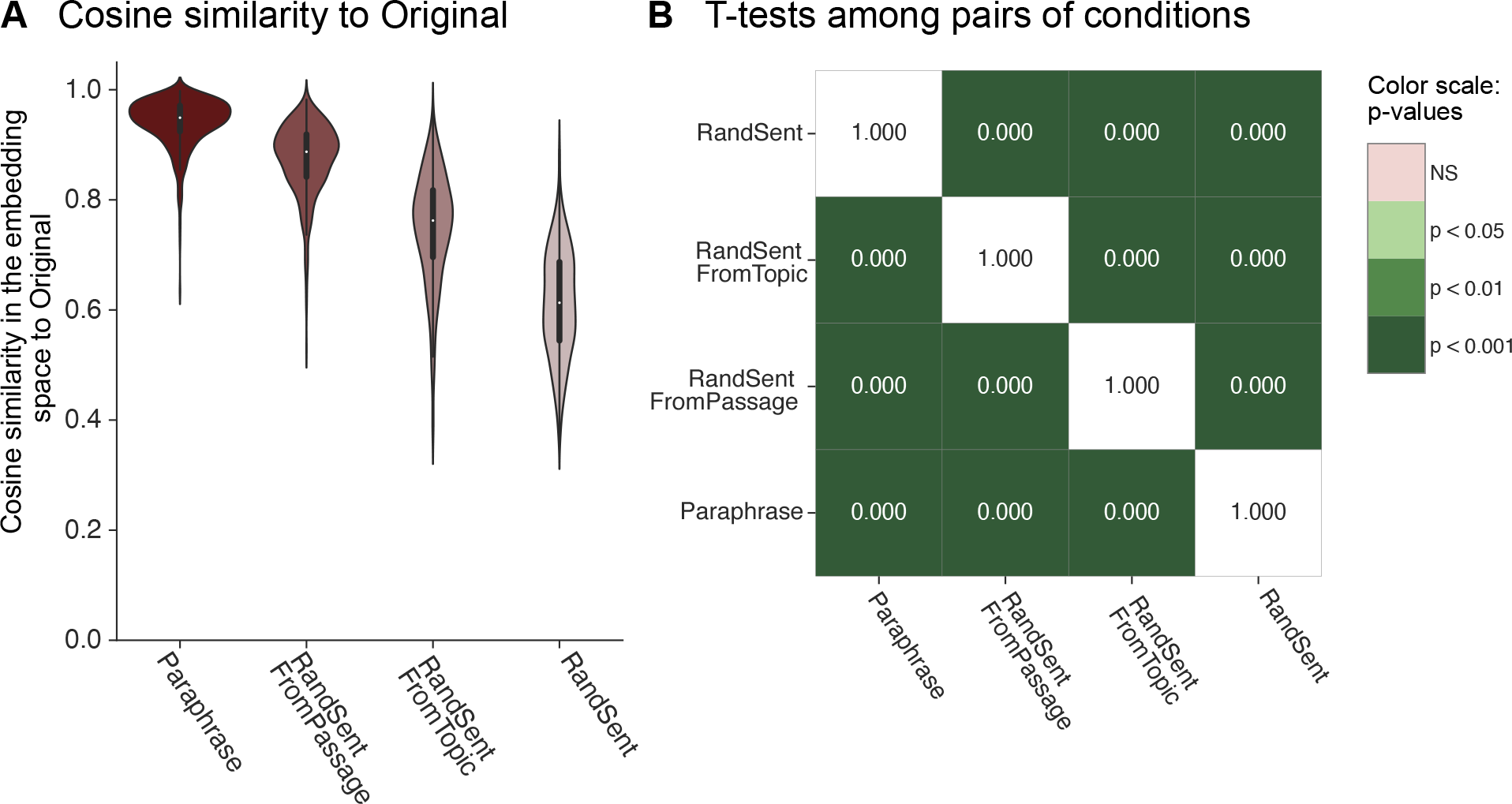
Validation of gradual semantic distance to Original within the semantic-distance perturbation conditions. (quantified using contextualized embeddings from GPT2-xl). **A.** We quantified the pairwise Cosine similarity of all 627 GPT2-xl sentence representation vectors for the *semantic-distance* manipulation datasets (conditions: *Paraphrase, RandSentFromPassage, RandSentFromTopic, RandSent*) with the representation of the intact version of the sentence (condition: *Original*). As expected, semantic similarity with the original sentence gradually decreased across conditions. **B.** The semantic-distance manipulations were significantly different from each other (independent two-sided t-tests with Bonferroni correction, ps<.001).

**Figure SI 5.**
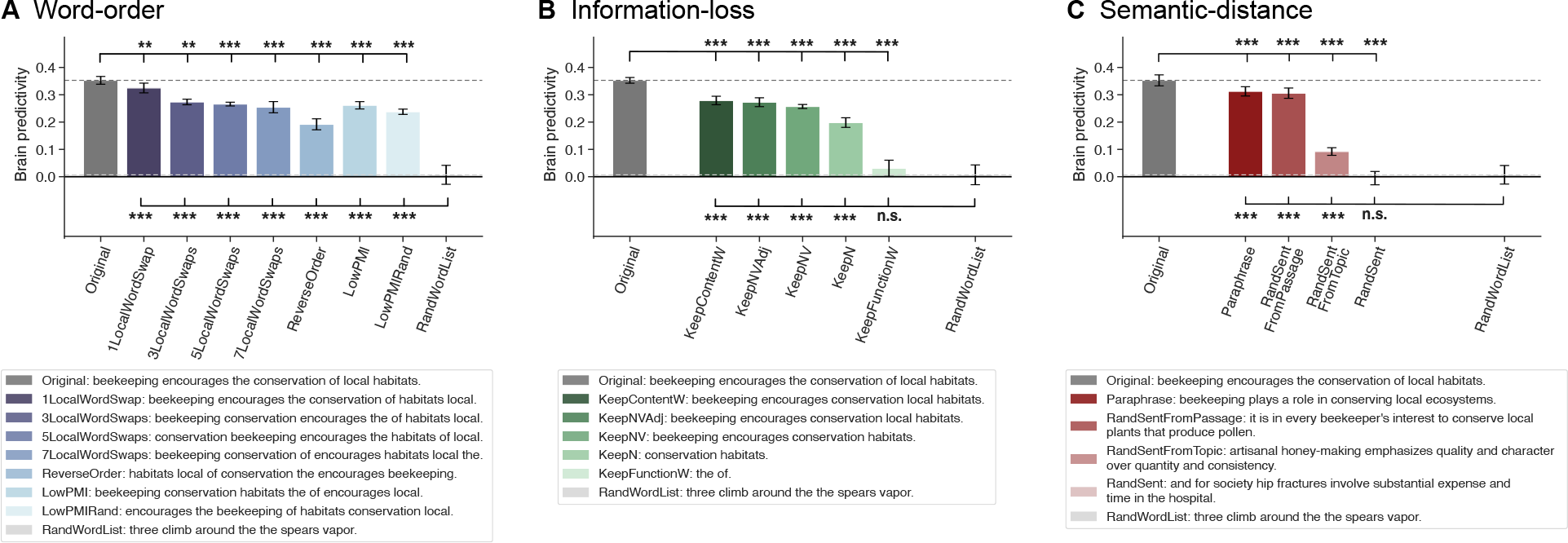
Brain predictivity performance of ANN-to-brain mapping models on held-out sentences using a fixed layer across all perturbation manipulation conditions. (as opposed to the best-performing layer for each condition as shown in Figure 2 in the main text). As in Figure 2, the mapping model was trained on ANN representations of intact sentences and evaluated on ANN representations of perturbed sentences (*TrainIntact-TestPerturbed_Contextualized*). Different from Figure 2, for this analysis, we selected the layer that performed best on the *Original* benchmark (encoder layer 44) instead of selecting the best-performing layer per condition. Bars show the brain predictivity using this fixed layer across the three perturbation manipulation conditions. Note that the *RandWordList* control condition reaches chance-level performance (zero).

**Figure SI 6.**
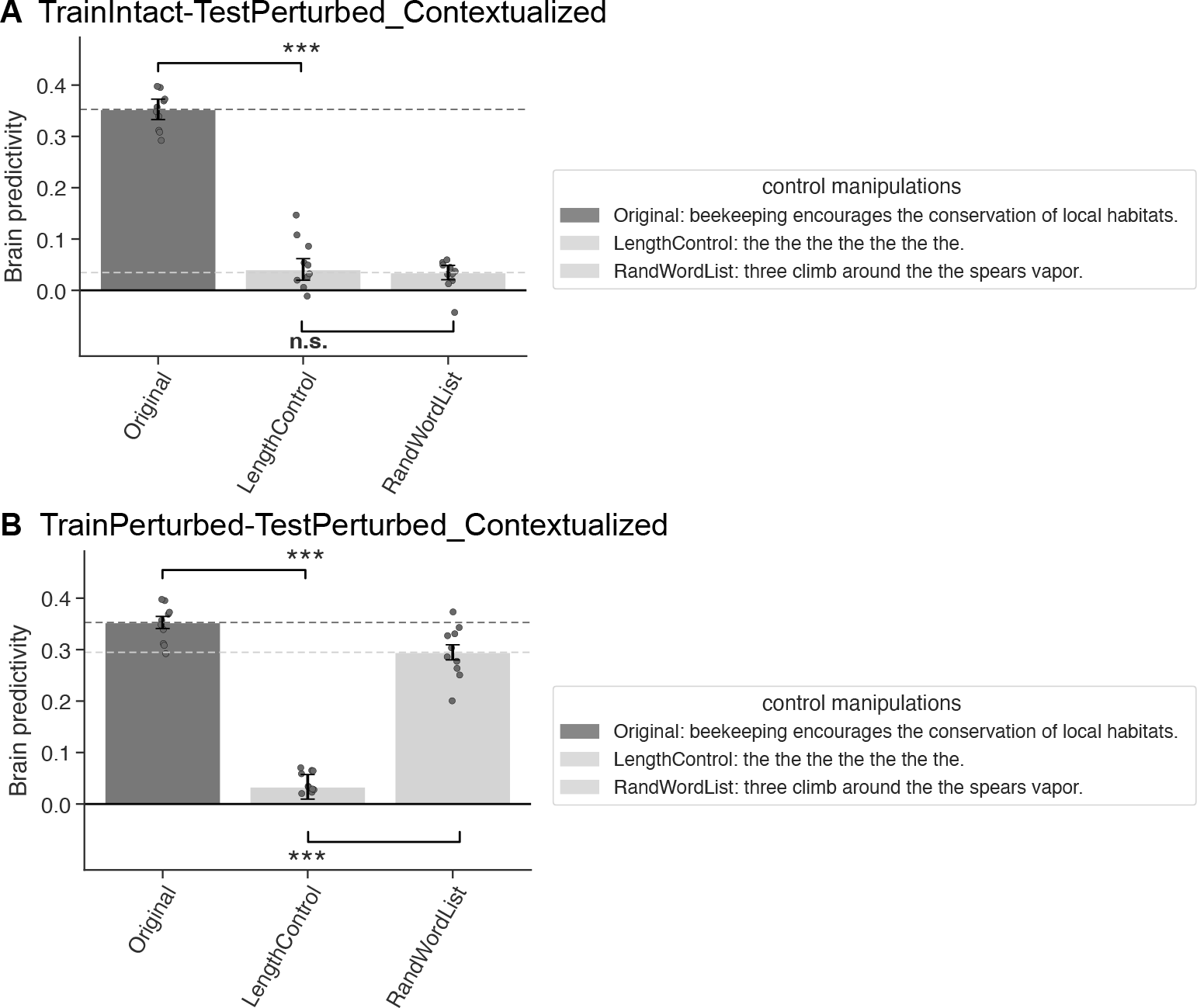
A length-controlled benchmark (*LengthControl*) performs on par with random word lists (*RandWordList*), and performs significantly better than chance. **A.** Performance of the ANN-to-brain mapping model on held-out sentences, trained on ANN representations of intact sentences and evaluated on ANN representations of perturbed sentences (*TrainIntact-TestPerturbed_Contextualized*) on an additional control condition *LengthControl* (each word in the original sentence replaced by the word “the”, allowing to test for effects due to the number of words in the sentence), relative to *Original* and *RandWordList*. A one-sample t-test shows that the *LengthControl* condition leads to non-zero predictivity performance (t=3.34, p<.01). **B.** Performance of ANN-to-brain mapping model on held-out sentences, trained on ANN representations of perturbed sentences and evaluated on ANN representations of perturbed sentences (*TrainPerturbed- TestPerturbed_Contextualized*), including the control condition *LengthControl*. A one-sample t-test shows that the *LengthControl* condition leads to non-zero predictivity performance (t=6.84, p<.001).

**Figure SI 7.**
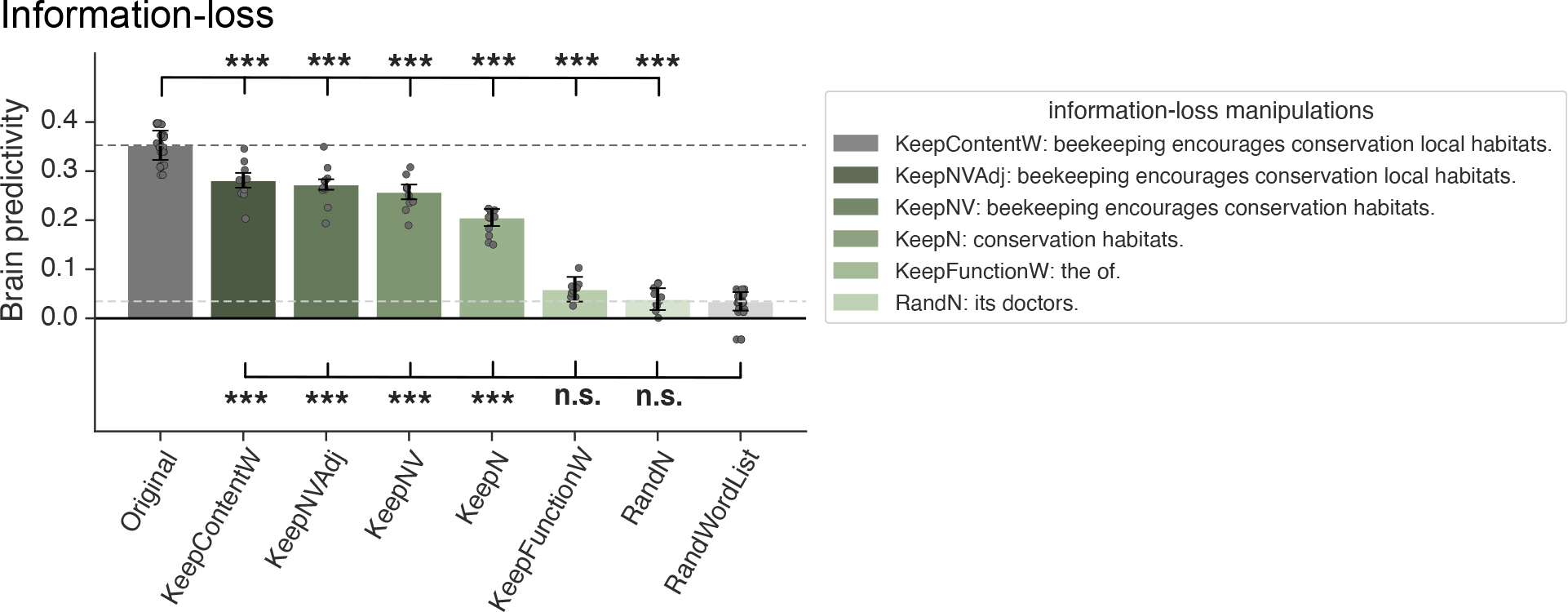
A benchmark with random nouns (*RandN*) performs on par with random word lists (*RandWordList*). The additional control benchmark, *RandN*, contained exclusively nouns (randomly sampled from the nouns in the dataset) and was matched for length with the KeepN condition. Brain predictivity performance of ANN-to-brain mapping model on held-out sentences (*TrainPerturbed-TestPerturbed_Contextualized*) of the *RandN* benchmark along with the remaining information-loss manipulations. The *RandN* benchmark performed on par with the random word list benchmark, *RandWordList*.

**Figure SI 8.**
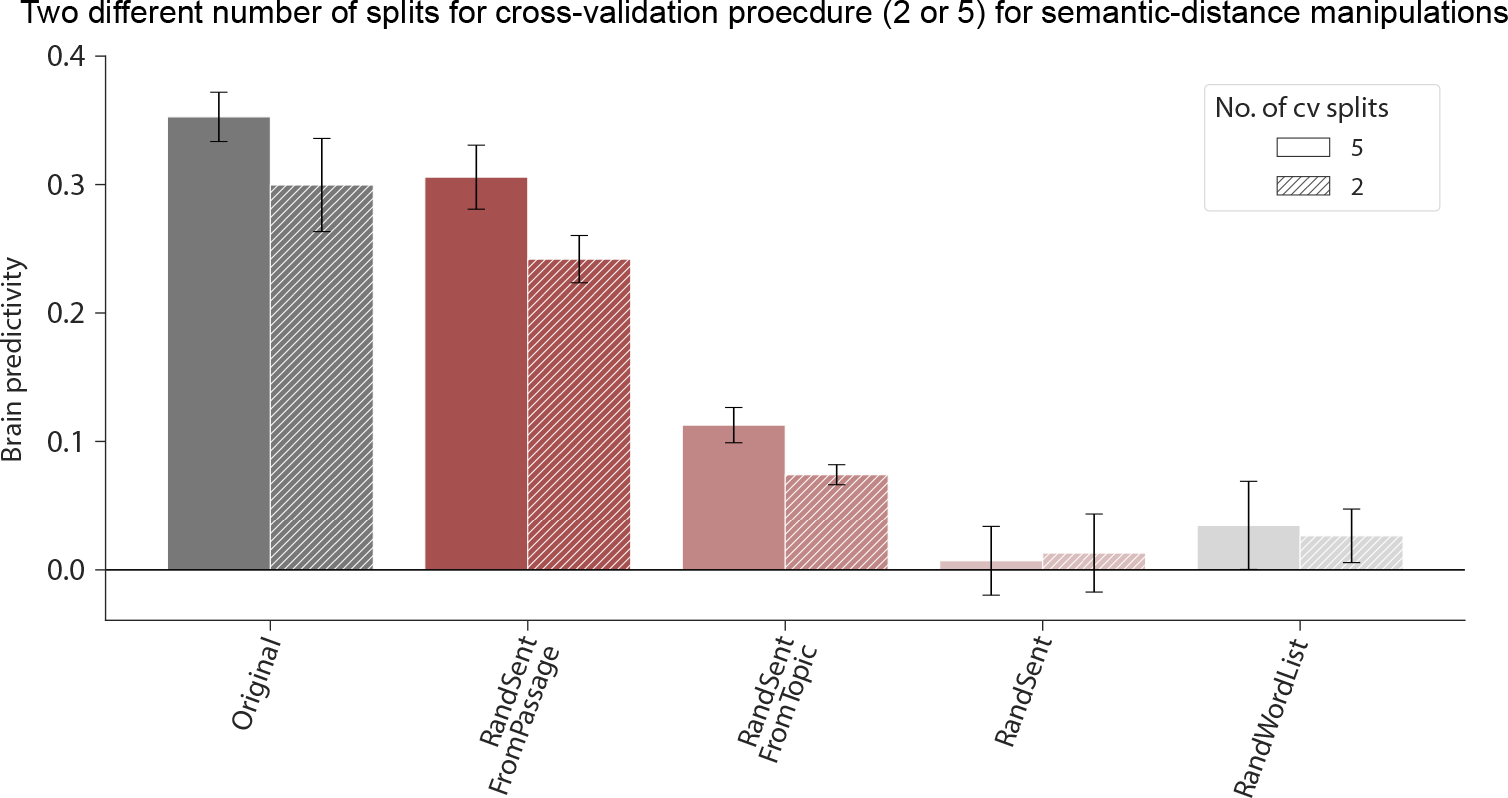
Key patterns of results are not affected by choice of number of splits in cross-validation procedure. Brain predictivity performance of ANN-to-brain mapping model on held-out sentences (*TrainIntact- TestPerturbed_Contextualized*) of the semantic-distance perturbation manipulation category benchmarks which were designed to shuffle sentences within the dataset (see <u>Methods; Perturbation manipulation conditions; Semantic-</u> <u>distance manipulations).</u> Due to the infeasibility of assigning sentences to 5 cross-validation folds and shuffling sentences according to the hierarchical structure of the Pereira et al. (2018) dataset, we additionally ran the *RandSent, RandSentFromPassage and RandSentFromTopic TrainIntact-TestPerturbed* benchmark versions (along with the *Original* and *RandWordList* benchmarks for comparison) using only 2 cross-validation splits instead of the default number of 5-folds. Using this procedure, all but 17.17% of sentence representations could be shuffled relative to its associated fMRI data for *RandSentFromPassage* and all sentences could be successfully shuffled with the associated fMRI data for *RandSentFromTopic*, leading to a less biased benchmark compared to the default 5-fold cross-validation scheme. The key patterns of results were not affected. For consistency with the remaining results, the 5-fold cross-validation results are reported in the main text.

**Figure SI 9.**
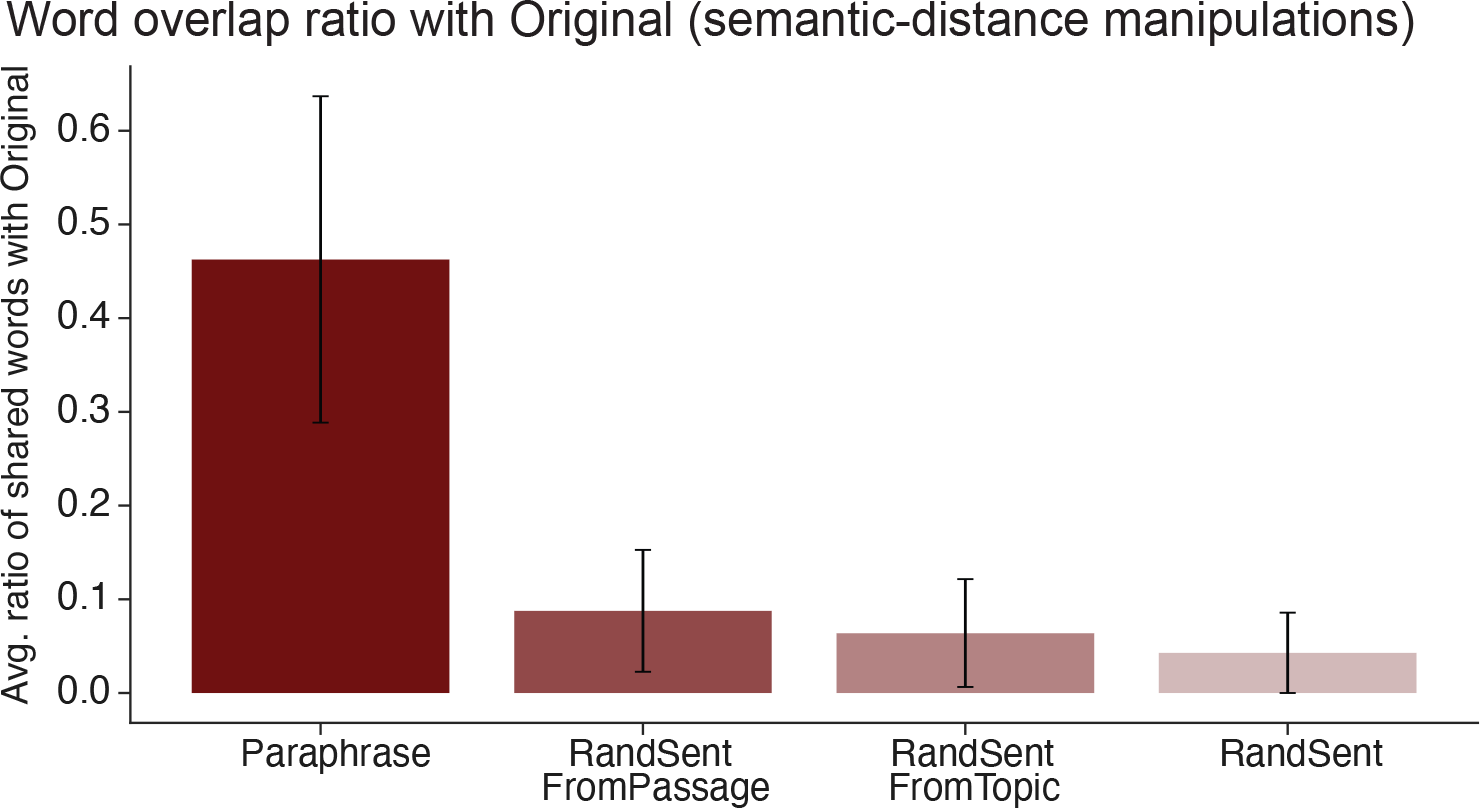
Quantification of word overlap ratio with *Original* across semantic-distance manipulations. For each of the semantic-distance manipulations, we quantified the ratio of word overlap for each sentence with the corresponding sentence from the *Original* condition and averaged the overlap ratios to obtain a summary statistic. Word overlap was quantified as (unique number of overlapping words) / (unique number of words in ({*semantic-distance manipulation*} + *Original*)). Error bars show the standard deviation from the mean.

**Figure SI 10.**
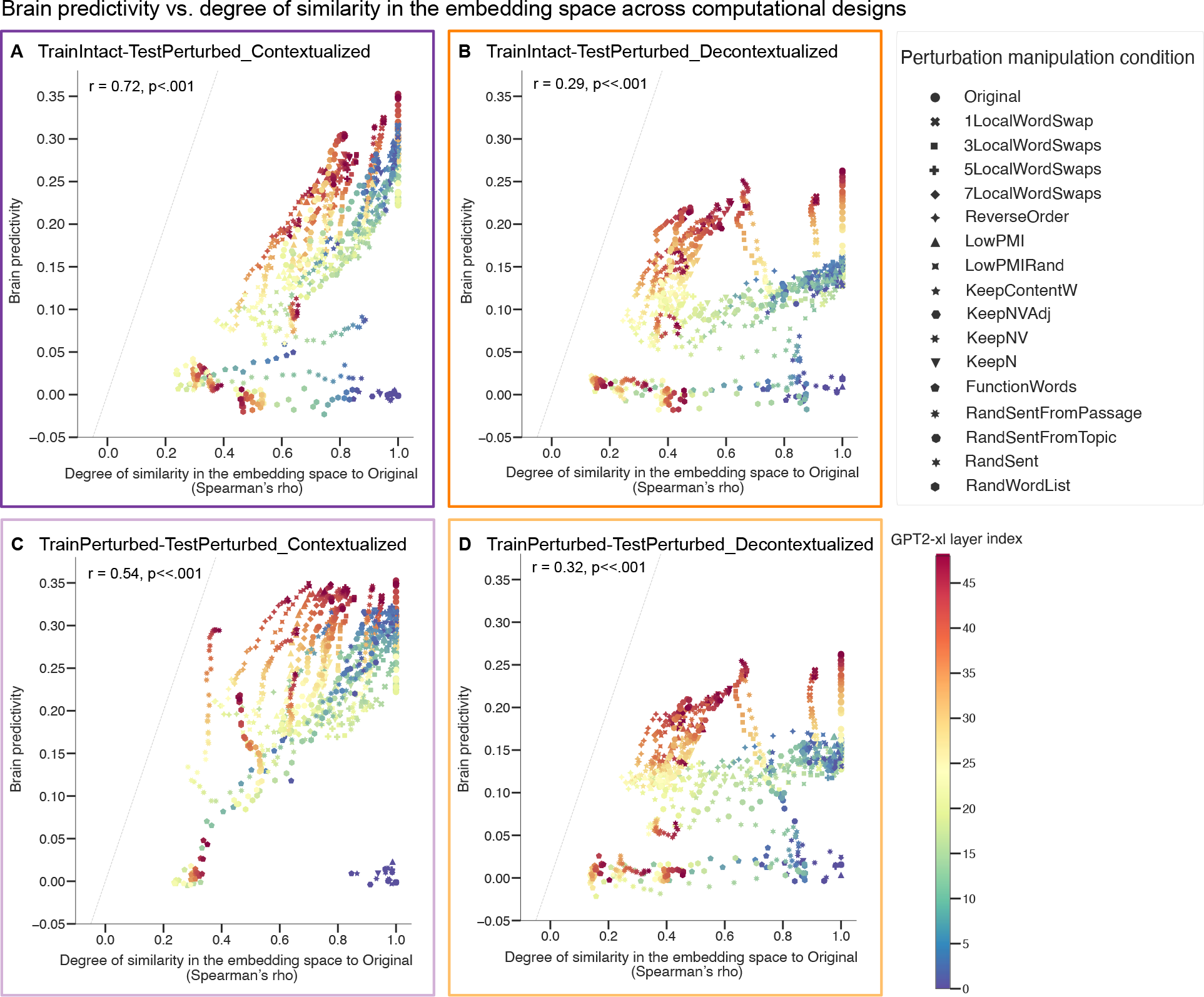
Representational similarity to the original sentences is correlated with brain predictivity. (across all computational experimental designs). Each individual data point shows the correlation between brain predictivity (y-axis) and degree of similarity to the intact sentence set (x-axis, quantified using the Spearman’s rank correlation coefficient, ρ) for a layer of the GPT2-xl ANN model and a certain perturbation manipulation condition for all computational experimental designs (panels A-D). The ANN layer index is denoted by colors. The perturbation manipulation condition is denoted by data point marker symbols.

**Figure SI 11.**
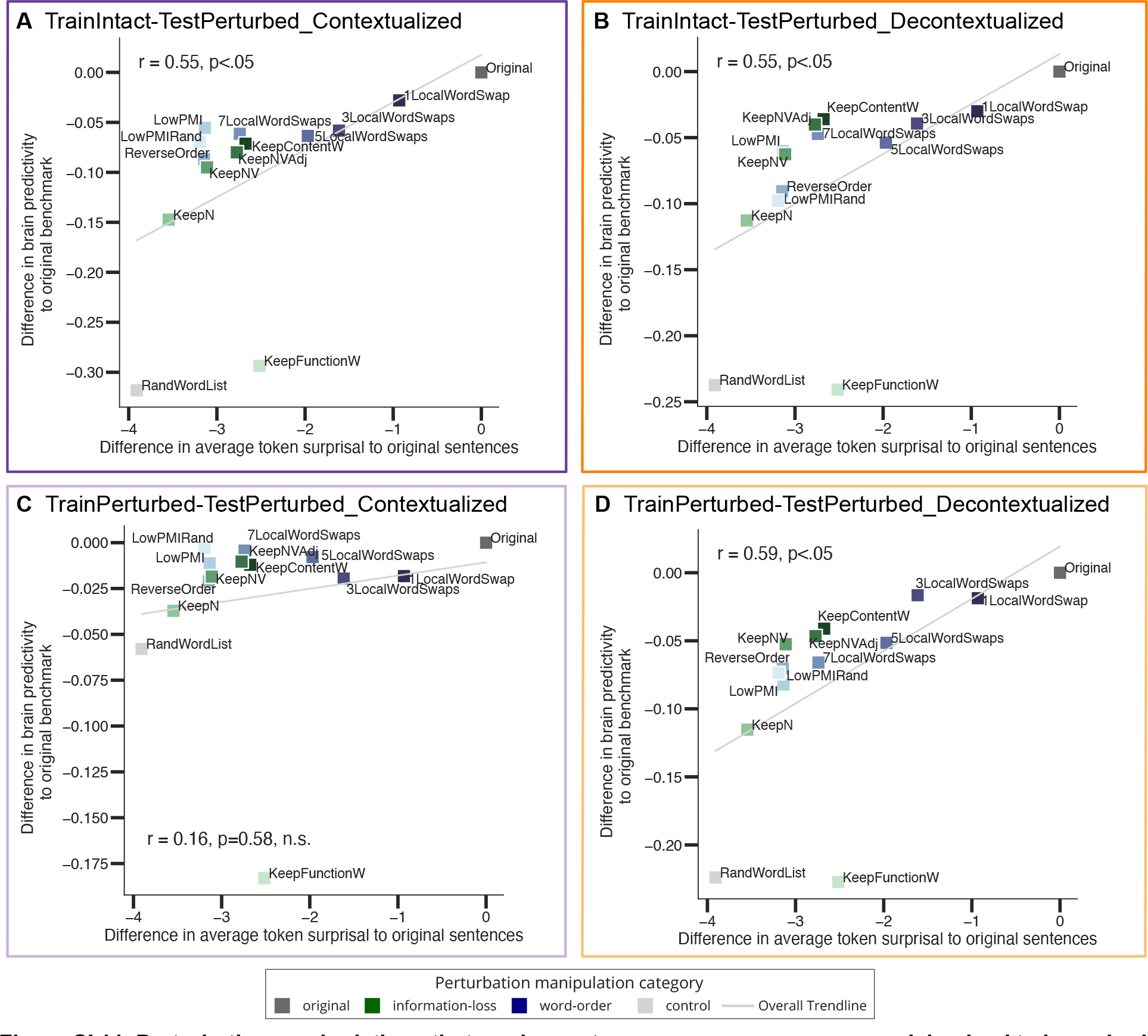
Perturbation manipulations that render sentences on average more surprising lead to lower brain predictivity. (across all computational experimental designs). The plots show the correlation between i) the difference in average sentence token surprisal between each perturbed sentence set and the original sentence set and ii) the difference in brain predictivity scores between each perturbed benchmark and the *Original* benchmark across all computational experimental designs (panels A-D).

**Figure SI 12.**
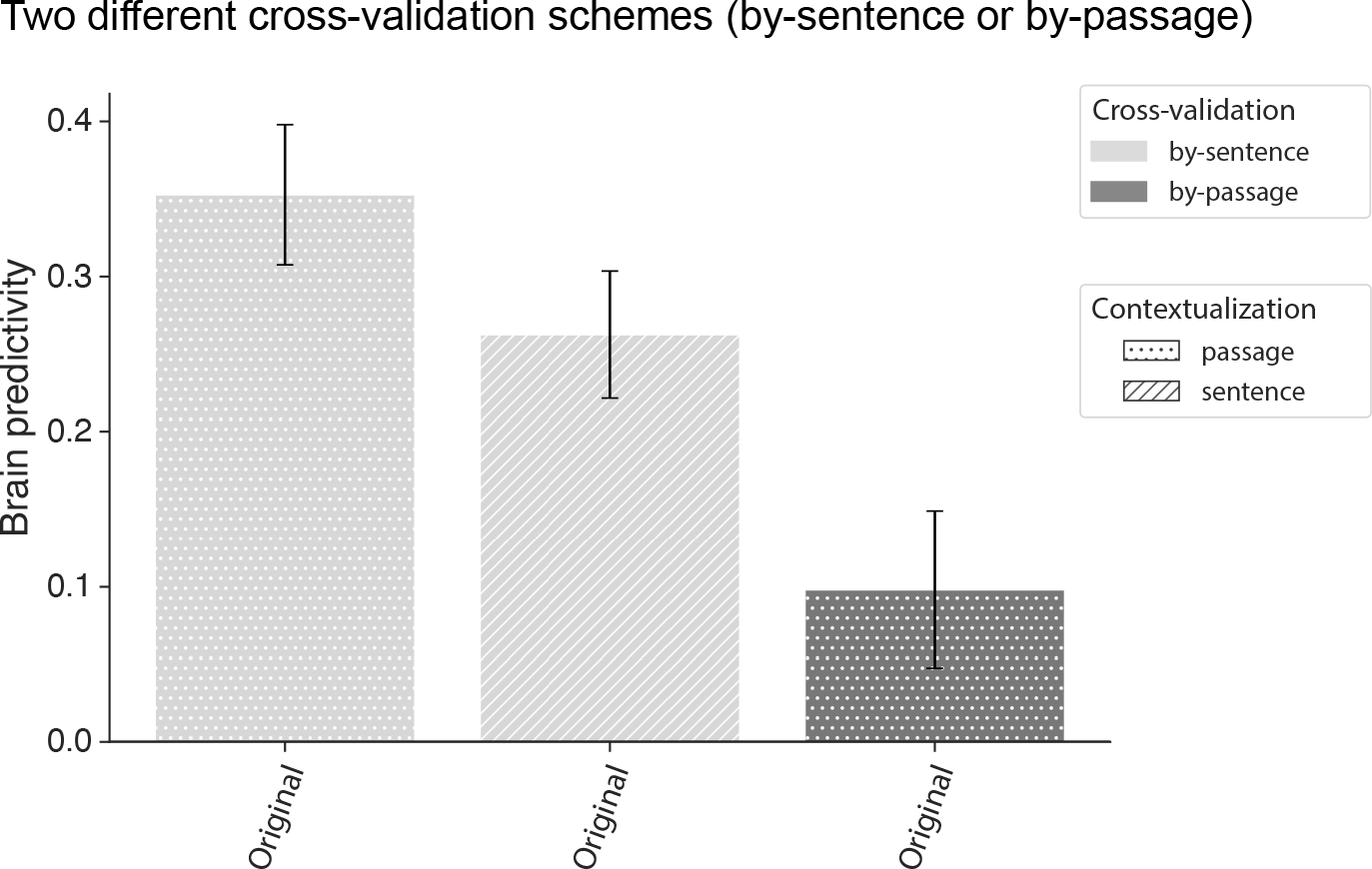
Brain predictivity performance of ANN-to-brain mapping using two different cross-validation schemes. Stimuli were separated into train and test sets in two ways: i) by sentence, disregarding the passage structure in the Pereira et al. 2018 dataset (light gray bars) vs. ii) by passage, such that all the sentences from the same passage end up in the same set (train or test) rather than being split across those sets (dark gray bar). For comparison to <u>Results;</u>Section 3.3, we show the contextualized and decontextualized ANN representation in the by-sentence split scheme as investigated throughout the manuscript. The ANN-to-brain mapping model was canonically trained (i.e., trained on the *Original,* intact sentences and tested on intact sentences (*TrainPerturbed-TestPerturbed_Contextualized*), consistent with prior work). Error bars show median absolute deviation (m.a.d.) error within participants.

### Supplementary Tables

**Table SI 1.**
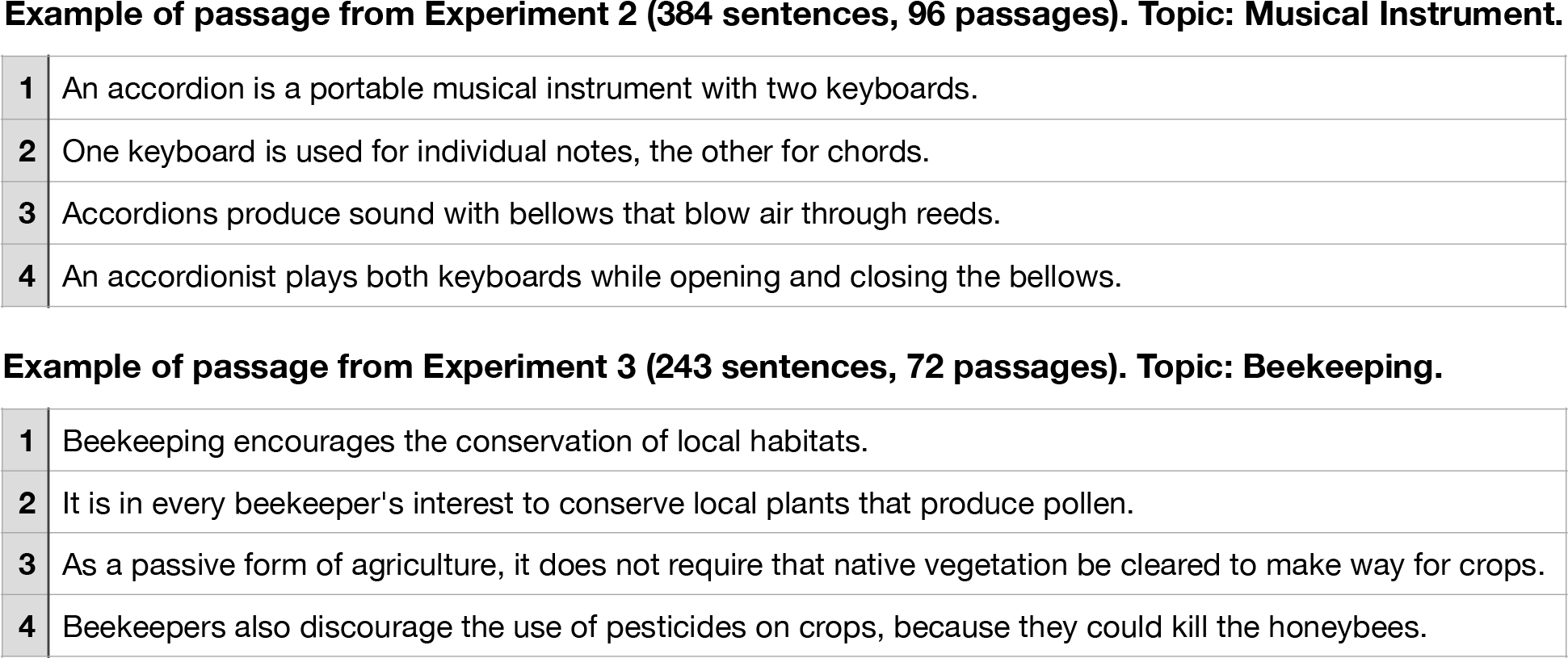
Examples of passages from Experiment 2 and Experiment 3 from Pereira et al. (2018), respectively.

**Example of passage from Experiment 2 (384 sentences, 96 passages). Topic: Musical Instrument.**
|  |  |
| --- | --- |
| 1 | An accordion is a portable musical instrument with two keyboards. |
| 2 | One keyboard is used for individual notes, the other for chords. |
| 3 | Accordions produce sound with bellows that blow air through reeds. |
| 4 | An accordionist plays both keyboards while opening and closing the bellows. |
| **Example of passage from Experiment 3 (243 sentences, 72 passages). Topic: Beekeeping.** |  |
| 1 | Beekeeping encourages the conservation of local habitats. |
| 2 | It is in every beekeeper's interest to conserve local plants that produce pollen. |
| 3 | As a passive form of agriculture, it does not require that native vegetation be cleared to make way for crops. |
| 4 | Beekeepers also discourage the use of pesticides on crops, because they could kill the honeybees. |

**Table SI 2.** **Statistics for Figure 2** in the main text (<u>Results;</u> Section 3.1). Pairwise, two-sided, dependent t-tests for all comparisons performed between the condition *Original* and all conditions of interest, as well as between condition *RandWordList* and all conditions of interest across perturbation manipulation classes. P-values were corrected for multiple comparisons (within each perturbation manipulation condition) using the Bonferroni procedure. Effect sizes, as quantified by Cohen’s d are reported.

**Table SI 1.** Examples of passages from Experiment 2 and Experiment 3 from Pereira et al. (2018), respectively.
|  | Comparison with *Original* |  |  | Comparison with *RandWordList* |  |  |
| --- | --- | --- | --- | --- | --- | --- |
|  | T-test statistic | Adjusted p-value | Cohen's d | T-test statistic | Adjusted p-value | Cohen's d |
| **1LocalWordSwap** | 5.517 | 0.0026 | 0.572 | -18.327 | 0.0000 | -8.288 |
| **3LocalWordSwaps** | 5.814 | 0.0018 | 1.527 | -17.437 | 0.0000 | -9.487 |
| **5LocalWordSwaps** | 6.507 | 0.0008 | 1.630 | -17.104 | 0.0000 | -9.322 |
| **7LocalWordSwaps** | 6.781 | 0.0006 | 1.641 | -16.790 | 0.0000 | -9.230 |
| **ReverseOrder** | 9.285 | 0.0000 | 2.383 | -15.265 | 0.0000 | -8.060 |
| **LowPMI** | 6.354 | 0.0009 | 1.560 | -20.505 | 0.0000 | -9.994 |
| **LowPMIRand** | 8.467 | 0.0001 | 2.043 | -17.035 | 0.0000 | -8.542 |
| **KeepContentW** | 9.209 | 0.0000 | 1.855 | -19.649 | 0.0000 | -7.141 |
| **KeepNVAAdj** | 12.825 | 0.0000 | 1.977 | -16.826 | 0.0000 | -6.617 |
| **KeepNV** | 17.944 | 0.0000 | 2.697 | -16.811 | 0.0000 | -6.942 |
| **KeepN** | 28.583 | 0.0000 | 4.842 | -13.441 | 0.0000 | -5.743 |
| **KeepFunctionW** | 17.400 | 0.0000 | 9.866 | -2.571 | 0.1506 | -1.122 |
| **Paraphrase** | 10.782 | 0.0000 | 1.238 | -17.966 | 0.0000 | -8.251 |
| **RandSentFromPassage** | 6.843 | 0.0003 | 0.975 | -17.600 | 0.0000 | -8.621 |
| **RandSentFromTopic** | 28.347 | 0.0000 | 7.502 | -6.438 | 0.0005 | -2.654 |
| **RandSent** | 24.262 | 0.0000 | 11.849 | 2.088 | 0.2657 | 1.002 |

**Table SI 3.** **Additional Statistics for Figure 2 in the main text (<u>Results;</u> Section 3.1).** Pairwise, two-sided, dependent t-tests for all comparisons performed between the conditions within the same perturbation manipulation classes. P-values were corrected for multiple comparisons (within each perturbation manipulation condition) using the Bonferroni procedure. Effect sizes, as quantified by Cohen’s d, are reported.

| Condition 1 | Condition 2 | T-test statistic | Adjusted p-value | Significance | Cohen's d | Manipulation |
| --- | --- | --- | --- | --- | --- | --- |
| 1LocalWordSwap | 3LocalWordSwaps | 2.9 | 0.369 | n.s. | 0.77 | word-order |
| 1LocalWordSwap | 5LocalWordSwaps | 3.469 | 0.148 | n.s. | 0.867 | word-order |
| 1LocalWordSwap | 7LocalWordSwaps | 3.437 | 0.156 | n.s. | 0.882 | word-order |
| 1LocalWordSwap | ReverseOrder | 6.886 | 0.002 | ** | 1.588 | word-order |
| 1LocalWordSwap | LowPMI | 3.568 | 0.127 | n.s. | 0.771 | word-order |
| 1LocalWordSwap | LowPMIRand | 5.775 | 0.006 | ** | 1.267 | word-order |
| 3LocalWordSwaps | 5LocalWordSwaps | 3.3 | 0.194 | n.s. | 0.127 | word-order |
| 3LocalWordSwaps | 7LocalWordSwaps | 3.076 | 0.278 | n.s. | 0.15 | word-order |
| 3LocalWordSwaps | ReverseOrder | 6.27 | 0.003 | ** | 1.055 | word-order |
| 3LocalWordSwaps | LowPMI | -0.161 | 1.0 | n.s. | -0.032 | word-order |
| 3LocalWordSwaps | LowPMIRand | 3.849 | 0.082 | n.s. | 0.648 | word-order |
| 5LocalWordSwaps | 7LocalWordSwaps | 0.403 | 1.0 | n.s. | 0.024 | word-order |
| 5LocalWordSwaps | ReverseOrder | 6.046 | 0.004 | ** | 0.929 | word-order |
| 5LocalWordSwaps | LowPMI | -0.898 | 1.0 | n.s. | -0.165 | word-order |
| 5LocalWordSwaps | LowPMIRand | 3.185 | 0.233 | n.s. | 0.523 | word-order |
| 7LocalWordSwaps | ReverseOrder | 5.07 | 0.014 | * | 0.9 | word-order |
| 7LocalWordSwaps | LowPMI | -0.927 | 1.0 | n.s. | -0.189 | word-order |
| 7LocalWordSwaps | LowPMIRand | 2.599 | 0.604 | n.s. | 0.496 | word-order |
| ReverseOrder | LowPMI | -7.609 | 0.001 | ** | -1.137 | word-order |
| ReverseOrder | LowPMIRand | -2.491 | 0.721 | n.s. | -0.4 | word-order |
| LowPMI | LowPMIRand | 3.82 | 0.086 | n.s. | 0.711 | word-order |
| Condition 1 | Condition 2 | T-test statistic | Adjusted p-value | Significance | Cohen's d | Manipulation |
| KeepContentW | KeepNVAdj | 1.224 | 1.0 | n.s. | 0.185 | information-loss |
| KeepContentW | KeepNV | 5.581 | 0.003 | ** | 0.684 | information-loss |
| KeepContentW | KeepN | 9.939 | 0.0 | *** | 2.516 | information-loss |
| KeepContentW | KeepFunctionW | 13.773 | 0.0 | *** | 7.019 | information-loss |
| KeepNVAdj | KeepNV | 3.167 | 0.114 | n.s. | 0.461 | information-loss |
| KeepNVAdj | KeepN | 10.063 | 0.0 | *** | 2.186 | information-loss |
| KeepNVAdj | KeepFunctionW | 11.576 | 0.0 | *** | 6.413 | information-loss |
| KeepNV | KeepN | 9.472 | 0.0 | *** | 1.923 | information-loss |
| KeepNV | KeepFunctionW | 12.889 | 0.0 | *** | 6.861 | information-loss |
| KeepN | KeepFunctionW | 9.804 | 0.0 | *** | 5.597 | information-loss |
| Condition 1 | Condition 2 | T-test statistic | Adjusted p-value | Significance | Cohen's d | Manipulation |
| Paraphrase | RandSentFromPassage | -1.476 | 1.0 | n.s. | -0.271 | semantic-distance |
| Paraphrase | RandSentFromTopic | 22.931 | 0.0 | *** | 6.089 | semantic-distance |
| Paraphrase | RandSent | 20.345 | 0.0 | *** | 10.255 | semantic-distance |
| RandSentFromPassage | RandSentFromTopic | 20.488 | 0.0 | *** | 6.45 | semantic-distance |
| RandSentFromPassage | RandSent | 24.504 | 0.0 | *** | 10.713 | semantic-distance |
| RandSentFromTopic | RandSent | 8.25 | 0.0 | *** | 4.231 | semantic-distance |

**Table SI 4.** **Overview of average sentence length across information-loss manipulation conditions** (*Original* condition included for reference). Note that the conditions in all other perturbation manipulation classes have the same number of words as the Original condition.

| Manipulation type | Condition | Average length<br>(in words) |
| --- | --- | --- |
| *Original* | *Original* | 12.33 |
| *Information-loss* | *KeepContentW* | 8.20 |
|  | *KeepNVAAdj* | 7.79 |
|  | *KeepNV* | 6.70 |
|  | *KeepN* | 4.35 |
|  | *RandN* | 4.35 |
|  | *KeepFunctionW* | 4.13 |

**Table SI 5.**
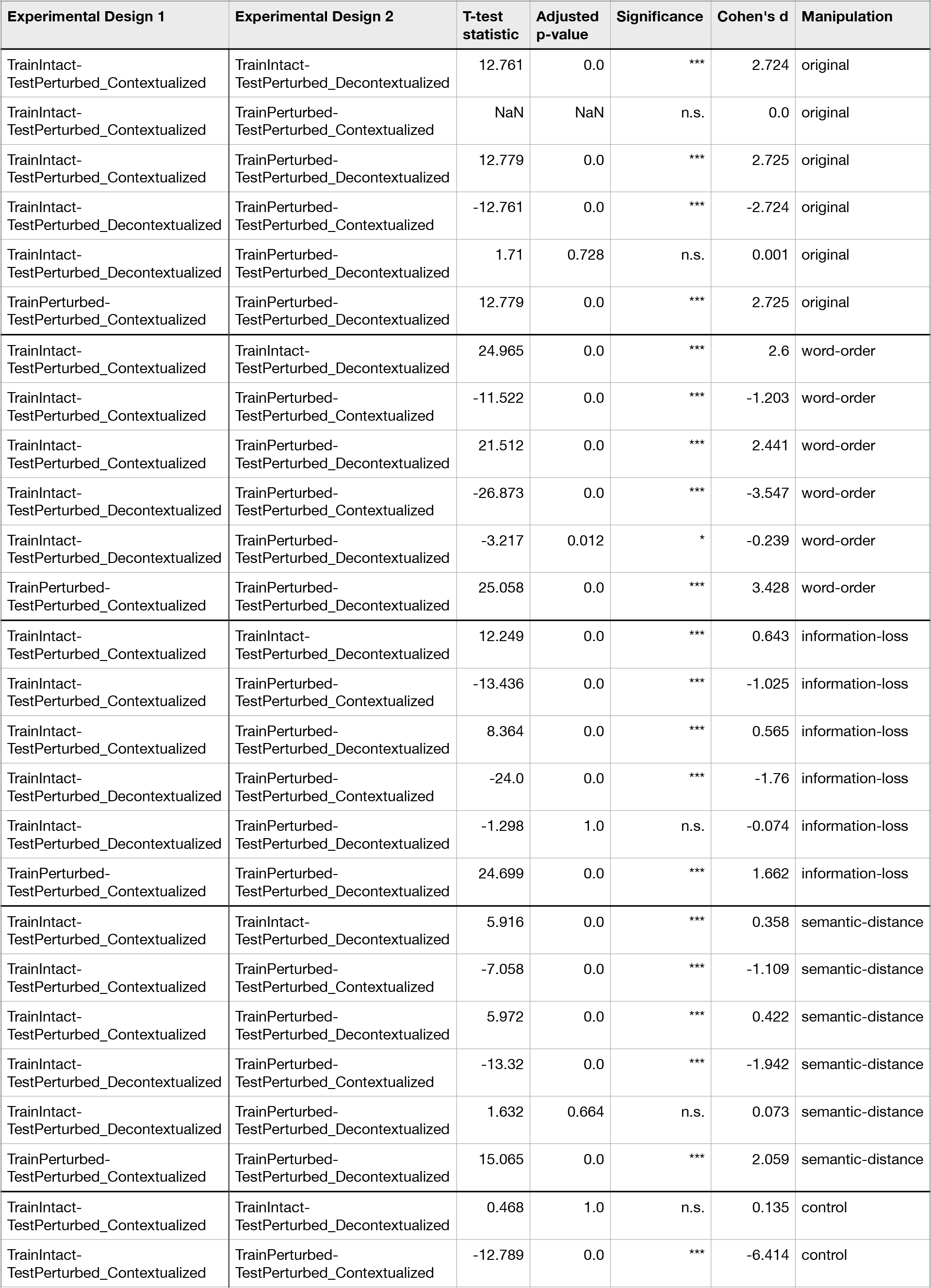
**Statistics for Figure 4 in the main text (<u>Results;</u> Section 3.3).** Pairwise, two-sided, dependent t-tests for all comparisons performed between the computational experimental design conditions across perturbation manipulation classes. P-values were corrected for multiple comparisons (within each perturbation manipulation condition) using the Bonferroni procedure.

| Experimental Design 1 | Experimental Design 2 | T-test statistic | Adjusted p-value | Significance | Cohen's d | Manipulation |
| --- | --- | --- | --- | --- | --- | --- |
| TrainIntact-TestPerturbed_Contextualized | TrainIntact-TestPerturbed_Decontextualized | 12.761 | 0.0 | *** | 2.724 | original |
| TrainIntact-TestPerturbed_Contextualized | TrainPerturbed-TestPerturbed_Contextualized | NaN | NaN | n.s. | 0.0 | original |
| TrainIntact-TestPerturbed_Contextualized | TrainPerturbed-TestPerturbed_Decontextualized | 12.779 | 0.0 | *** | 2.725 | original |
| TrainIntact-TestPerturbed_Decontextualized | TrainPerturbed-TestPerturbed_Contextualized | -12.761 | 0.0 | *** | -2.724 | original |
| TrainIntact-TestPerturbed_Decontextualized | TrainPerturbed-TestPerturbed_Decontextualized | 1.71 | 0.728 | n.s. | 0.001 | original |
| TrainPerturbed-TestPerturbed_Contextualized | TrainPerturbed-TestPerturbed_Decontextualized | 12.779 | 0.0 | *** | 2.725 | original |
| TrainIntact-TestPerturbed_Contextualized | TrainIntact-TestPerturbed_Decontextualized | 24.965 | 0.0 | *** | 2.6 | word-order |
| TrainIntact-TestPerturbed_Contextualized | TrainPerturbed-TestPerturbed_Contextualized | -11.522 | 0.0 | *** | -1.203 | word-order |
| TrainIntact-TestPerturbed_Contextualized | TrainPerturbed-TestPerturbed_Decontextualized | 21.512 | 0.0 | *** | 2.441 | word-order |
| TrainIntact-TestPerturbed_Decontextualized | TrainPerturbed-TestPerturbed_Contextualized | -26.873 | 0.0 | *** | -3.547 | word-order |
| TrainIntact-TestPerturbed_Decontextualized | TrainPerturbed-TestPerturbed_Decontextualized | -3.217 | 0.012 | * | -0.239 | word-order |
| TrainPerturbed-TestPerturbed_Contextualized | TrainPerturbed-TestPerturbed_Decontextualized | 25.058 | 0.0 | *** | 3.428 | word-order |
| TrainIntact-TestPerturbed_Contextualized | TrainIntact-TestPerturbed_Decontextualized | 12.249 | 0.0 | *** | 0.643 | information-loss |
| TrainIntact-TestPerturbed_Contextualized | TrainPerturbed-TestPerturbed_Contextualized | -13.436 | 0.0 | *** | -1.025 | information-loss |
| TrainIntact-TestPerturbed_Contextualized | TrainPerturbed-TestPerturbed_Decontextualized | 8.364 | 0.0 | *** | 0.565 | information-loss |
| TrainIntact-TestPerturbed_Decontextualized | TrainPerturbed-TestPerturbed_Contextualized | -24.0 | 0.0 | *** | -1.76 | information-loss |
| TrainIntact-TestPerturbed_Decontextualized | TrainPerturbed-TestPerturbed_Decontextualized | -1.298 | 1.0 | n.s. | -0.074 | information-loss |
| TrainPerturbed-TestPerturbed_Contextualized | TrainPerturbed-TestPerturbed_Decontextualized | 24.699 | 0.0 | *** | 1.662 | information-loss |
| TrainIntact-TestPerturbed_Contextualized | TrainIntact-TestPerturbed_Decontextualized | 5.916 | 0.0 | *** | 0.358 | semantic-distance |
| TrainIntact-TestPerturbed_Contextualized | TrainPerturbed-TestPerturbed_Contextualized | -7.058 | 0.0 | *** | -1.109 | semantic-distance |
| TrainIntact-TestPerturbed_Contextualized | TrainPerturbed-TestPerturbed_Decontextualized | 5.972 | 0.0 | *** | 0.422 | semantic-distance |
| TrainIntact-TestPerturbed_Decontextualized | TrainPerturbed-TestPerturbed_Contextualized | -13.32 | 0.0 | *** | -1.942 | semantic-distance |
| TrainIntact-TestPerturbed_Decontextualized | TrainPerturbed-TestPerturbed_Decontextualized | 1.632 | 0.664 | n.s. | 0.073 | semantic-distance |
| TrainPerturbed-TestPerturbed_Contextualized | TrainPerturbed-TestPerturbed_Decontextualized | 15.065 | 0.0 | *** | 2.059 | semantic-distance |
| TrainIntact-TestPerturbed_Contextualized | TrainIntact-TestPerturbed_Decontextualized | 0.468 | 1.0 | n.s. | 0.135 | control |
| TrainIntact-TestPerturbed_Contextualized | TrainPerturbed-TestPerturbed_Contextualized | -12.789 | 0.0 | *** | -6.414 | control |
| TrainIntact-TestPerturbed_Contextualized | TrainPerturbed-TestPerturbed_Decontextualized | -0.05 | 1.0 | n.s. | -0.014 | control |
| TrainIntact-TestPerturbed_Decontextualized | TrainPerturbed-TestPerturbed_Contextualized | -12.437 | 0.0 | *** | -6.767 | control |
| TrainIntact-TestPerturbed_Decontextualized | TrainPerturbed-TestPerturbed_Decontextualized | -0.519 | 1.0 | n.s. | -0.151 | control |
| TrainPerturbed-TestPerturbed_Contextualized | TrainPerturbed-TestPerturbed_Decontextualized | 13.808 | 0.0 | *** | 6.421 | control |

## Supplementary Methods

### Estimation of ceiling

Due to intrinsic noise in biological measurements, we estimated a ceiling value (i.e., a normalizing constant for the correlation between actual and ANN-predicted brain responses) which quantifies how well the best possible “average human” model could perform on predicting brain responses in single voxels for held-out “target” participants. In our ceiling estimation, we included the *n*=5 participants that completed both experiments in the Pereira et al. (2018) dataset to obtain full overlap in the materials across participants. Following Schrimpf et al. (2021), the ceiling value was estimated using a three-step procedure:

#### Step 1: Collect data for extrapolation

We first subsampled the data with *n*=5 recorded participants into all possible combinations of p participants for all *p*∈[2,n=5]. For example, for *p*=2 there are 5P2=20 possible subsample combinations from the participant pool. To keep computational cost manageable, for each subsample size *p*, we used only 10 of these random subsample combinations to calculate correlation scores (that will be used in the extrapolation in Step 2). For example, for p=2, we randomly picked 10 subsamples from the 10 possible participant subsamples of size 2 from the 5 participants in the participant pool. For each of these subsamples for a given *p*, we then designated one participant as the target (such that each participant is chosen as the target once) whose brain responses we attempt to predict from the remaining *p*-1 participants (e.g., predict 1 participant from 1 (other) participant (*p*=2), 1 from 2 participants (*p*=3), …, 1 from *n-1* participants (*p*=n)) to obtain a correlation score between the predicted and the actual activation of the voxel for each voxel in the “target” participant for the given subsample. Hence, instead of predicting a “target” participant’s voxels using an ANN embedding (as done in the main analysis), we used voxels from the “predictor” participants as an embedding (if only one participant was used as the “predictor” participant, we used that participant’s voxels; if two or more participants were used as “predictor” participants, their voxels were concatenated).

#### Step 2: Get ceiling value per voxel by extrapolating prediction accuracy to infinitely many participants

We computed a ceiling value for each voxel individually using an extrapolation approach. To extrapolate the approach described in Step 1 beyond the number of participants in the participant pool (*n*=5), we fitted the equation for each voxel where p is each subsample’s number of participants (i.e., 2, …, 5), v is each subsample’s correlation score and are the fitted parameters for asymptote and slope respectively. For each voxel, we used 100 bootstraps to fit the ceiling to different subsets of predicted values across subsamples and used the median of the asymptote values from the 100 bootstraps as that voxel’s ceiling value. Specifically, for each of the 100 bootstraps, we resample the correlation values (with replacement) for each *p* and fit the equation above.

#### Step 3: Aggregate the ceiling values across voxels to obtain the ceiling value

After estimating a ceiling value for all voxels in all of the 5 participants as described in Steps 1 and 2, we used these scores to compute the dataset’s final ceiling value that was used as a normalizing constant for the brain predictivity scores (correlation between actual and ANN-predicted brain responses). To do so, we computed the median of the per-voxel ceilings across all voxels and all participants. Via this procedure, we obtained a final ceiling value of 0.32 for the Pereira et al. (2018) dataset. The model scores we report are the model’s overall raw correlation scores (aggregated as described in <u>Methods; Comparison of ANN model representations to brain</u> <u>measurements</u>) divided by this ceiling value.

## Footnotes

1 We used positive pointwise mutual information because negative PMI values are in practice extremely noisy due to data sparsity (Jurafsky & Martin, 2008).

2 The contextualization for the test set sentences in the *TrainIntact-TestPerturbed_Contextualized* semantic-distance manipulation benchmarks where sentence representations were shuffled relative to the fMRI data (*RandSentFromPassage, RandSentFromTopic, RandSent*) is an exception to this rule. Here, we take the sentence representations of the original sentences and shuffle the order of these (correctly contextualized) sentence representations with the associated fMRI data either randomly or based on the sentence’s membership in a particular passage or topic.

